# Disrupted Autophagy and Neuronal Dysfunction in *C. elegans* Knock-in Models of FUS Amyotrophic Lateral Sclerosis

**DOI:** 10.1101/799932

**Authors:** Saba N. Baskoylu, Natalie Chapkis, Burak Unsal, Jeremy Lins, Kelsey Schuch, Jonah Simon, Anne C. Hart

## Abstract

It remains unclear how mutations in FUS, a ribonucleoprotein, lead to neuronal dysfunction in Amyotrophic Lateral Sclerosis (ALS) patients. To examine mechanisms underlying ALS FUS dysfunction, we generated the first *C. elegans* knock-in models using CRISPR/Cas9-mediated genome editing, creating R524S and P525L ALS FUS models. Although FUS inclusions were not detected, ALS FUS animals showed defective neuromuscular function, as well as stress-induced locomotion defects. Unlike *C. elegans* lacking the endogenous FUS ortholog, ALS FUS animals had impaired neuronal autophagy and increased SQST-1 accumulation in ALS FUS motor neurons. Loss of *sqst-1*, the *C. elegans* ortholog for ALS-linked, autophagy adaptor protein SQSTM1/p62, suppressed both neuromuscular and stress-induced locomotion defects in ALS FUS animals, but did not suppress neuronal autophagy defects. Therefore, autophagy dysfunction is upstream of, and not dependent on, SQSTM1 function in ALS FUS pathogenesis. Combined, our findings demonstrate that autophagy dysfunction likely contributes to protein homeostasis and neuromuscular defects in ALS FUS knock-in animals.

## Introduction

Amyotrophic Lateral Sclerosis (ALS) is an incurable adult-onset neurodegenerative disease characterized by progressive neuromuscular denervation and motor neuron loss. Mutations in FUsed in Sarcoma (FUS) account for approximately 5% and 1% of familial and sporadic ALS cases, respectively [1]. FUS is an DNA/RNA binding protein that participates in transcription, splicing and mRNA stabilization in the nucleus, and contributes to stress granule formation, axonal transport and microRNA function in the cytoplasm [2]. Most disease-linked FUS mutations map to the FUS C-terminal nuclear localization signal (NLS), leading to cytoplasmic FUS mislocalization and aggregation in patient motor neurons [3]. Cytoplasmic FUS aggregates are also seen in patients with Frontotemporal Dementia (FTD) [4]. FUS overexpression in mice leads to dose-dependent neuromuscular defects and neurodegeneration reminiscent of ALS, with altered RNA transcription and splicing in motor neurons [2]. FUS overexpression impairs synaptic function in mice, flies and nematodes [5–11]. Furthermore, in fly and mouse ALS FUS overexpression models, neuromuscular dysfunction precedes motor neuron degeneration [12,13], suggesting synaptic dysfunction is an early event in ALS pathogenesis. These over/mis-expression models demonstrate that FUS protein perturbation is deleterious for neuronal function and survival. However, molecular changes that occur before symptom onset remain elusive. ALS FUS knock-in model in mice showed neuronal dysfunction and degeneration in the absence of cytoplasmic FUS protein accumulation, suggesting aggregates may not be the initial drivers of neuronal dysfunction in neurons [14].

Studying early changes in ALS FUS may yield insight into disease mechanisms. Defective autophagy likely contributes to neuronal dysfunction in neurodegenerative diseases, as evidenced by ALS-linked mutations in autophagy-related genes SQSTM1/p62 [15], UBQLN2 [16], VCP [17], OPTN [12,18,19] and TBK1 [5]. Autophagy is a catabolic pathway that delivers membrane-enclosed proteins and organelles to lysosomes. Essential autophagy-mediated cellular functions include protein turnover, misfolded/aggregation-prone protein degradation, and metabolic stress adaptation [20]. Autophagy adaptor protein SQSTM1 accumulates in ALS patient motor neurons [21], which may indicate autophagy dysfunction. Consistent with this, LC3-positive autophagy vesicles are elevated in ALS FUS patient motor neurons [22,23]. Thus, autophagy dysfunction may contribute to ALS disease pathogenesis.

Studies in FUS overexpression models have revealed autophagy dysfunction. FUS overexpression perturbs autophagy flux in neuronal cell lines transfected with wild type or mutant FUS [11,23]. Wild type FUS overexpression alters RNA processing, upregulates lysosomal genes, and leads to improper SQSTM1 accumulation in transgenic mice [11]. Combined, these findings suggest that improper FUS levels negatively impact autophagy. However, it remains unclear if ALS-linked FUS mutations *per se* impact neuronal autophagy at early stages of pathogenesis.

Here, we generated the first ALS FUS knock-in models in *C. elegans*. Using CRISPR/Cas9-mediated homologous recombination, we inserted the equivalent patient mutations for R524S and P525L into the *C. elegans* FUS orthologous gene, *fust-1*. Animals carrying these knock-in alleles were hypersensitive to stress and had impaired neuromuscular function. Importantly, FUS knock-in mutations differentially impaired neuronal and muscular autophagy, while leaving the neuronal ubiquitin proteasome system intact. Loss of function in the SQSTM1 ortholog, *sqst-1*, suppressed stress-induced locomotion defects and aldicarb sensitivity in ALS FUS animals.. However, loss of SQSTM1/*sqst-1* function did not suppress neuronal autophagy defects in ALS FUS animals, despite the restoration of normal locomotion and aldicarb sensitivity. Our findings in new *C. elegans* ALS FUS knock-in models complement ALS FUS overexpression models and suggest autophagy dysfunction contributes to ALS FUS pathogenesis.

## Results

### ALS FUST-1 protein mislocalizes to the cytoplasm in *C. elegans* motor neurons

At steady state levels, wild type FUS protein localizes primarily to the nucleus. ALS-linked FUS mutations R524S and P525L disrupt nuclear FUS localization and lead to cytoplasmic FUS aggregation [22]. We examined whether the introduction of equivalent nuclear localization signal (NLS) mutations would disrupt subcellular localization of FUST-1 protein, the *C. elegans* ortholog for FUS (Figure 1A). The R524S and P525L equivalent point mutations in the *C. elegans* FUST-1 protein are R446S and P447L, respectively. Note that *C. elegans fust-1* alleles generated herein will be referred to with amino acid numbering in the orthologous human FUS protein.

**Figure 1.**
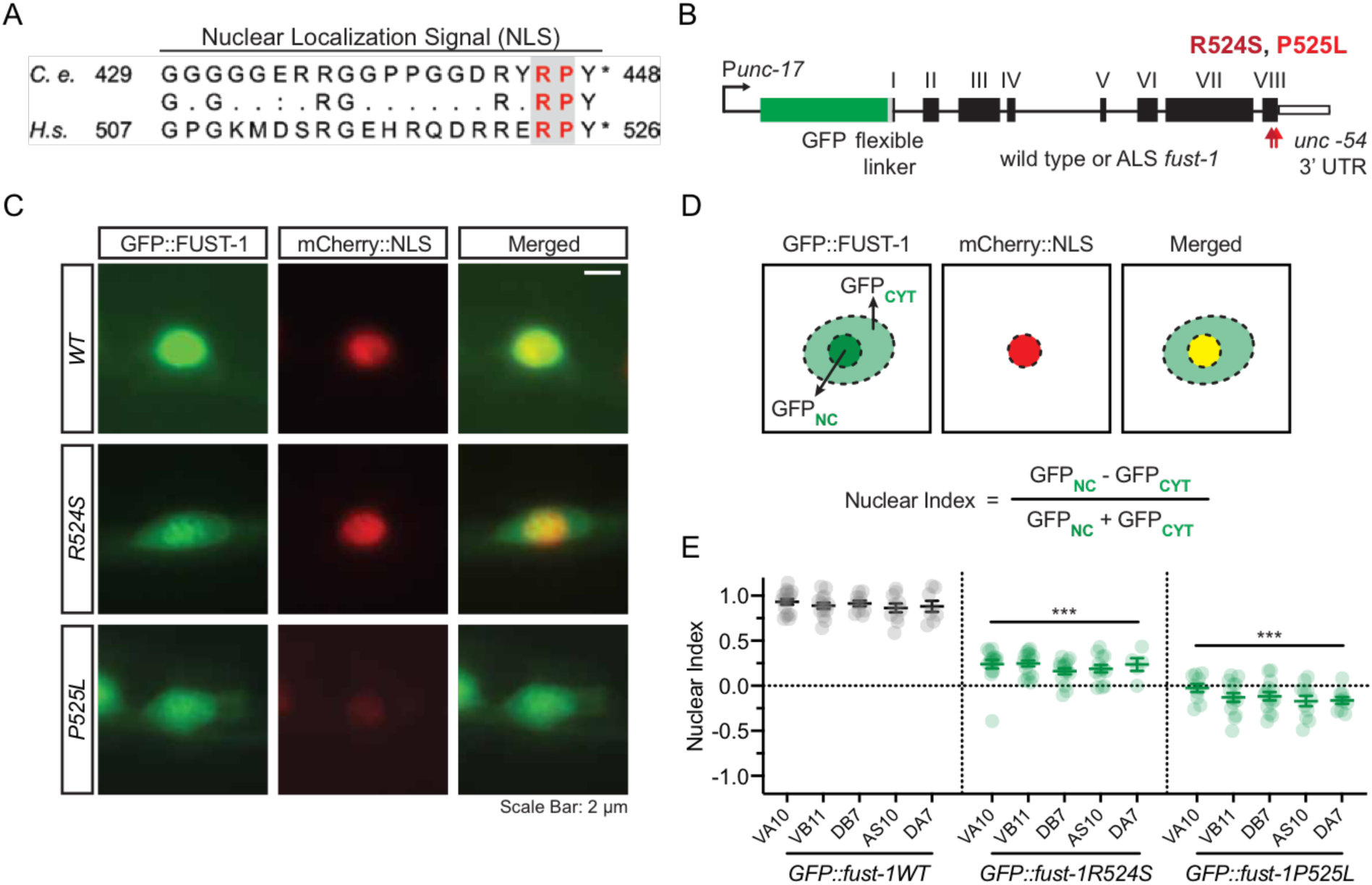
Cytoplasmic mislocalization of *C. elegans* FUST-1 containing R524S and P525L in motor neurons. **(A)** Alignment of C-terminal nuclear localization signal motif in *C. elegans* FUST-1 and human FUS proteins. Identical amino acid residues are indicated between the protein sequences; conserved and semi-conserved amino acid residues are indicated with “:” and “.”, respectively. Red residues were mutated herein to create *C. elegans* R524S and P525L ALS FUS models. *Homo sapiens* (*H.s.*) FUS GenPept Accession Number CAG33028.1; *C. elegans* (*C.e.*) FUST-1 GenPept Accession Number NP_495483.1. For alignment of complete human FUS and *C. elegans* FUST-1 protein sequences, see Supplementary Figure 1. **(B)** Transgenes used to determine relative nuclear-to-cytoplasmic GFP-tagged FUST-1 localization in cholinergic motor neurons using the *unc-17* cholinergic promoter (GFP::FUST-1). All constructs contain a flexible linker sequence between the N-terminal GFP fusion tag and the FUST-1 protein, and use the *unc-54* 3’ untranslated region (UTR). The black boxes show *fust-1* exons. Red arrows point at the location of ALS-associated FUST-1 mutations. **(C)** Representative images of wild type and mutant GFP::FUST-1 in motor neurons. The neuronal nuclei were labelled with mCherry::NLS driven under the *cho-1* cholinergic promoter. Scale bar = 2 um. **(D)** Illustration of the nuclear and cytoplasmic fluorescence associated with GFP::FUST-1 fluorescence in motor neurons. GFP fluorescence was measured in the nucleus and in the cytoplasm. To quantify the relative nuclear-to-cytoplasmic GFP::FUST-1 localization in motor neurons, neuronal index was determined as shown in the equation. A nuclear index of 1 corresponds to predominantly nuclear-localized protein, and a nuclear index of −1 corresponds to predominantly cytoplasmic-localized protein. **(E)** Nuclear index of the GFP::FUST-1 fluorescence was measured in five motor neurons along the ventral nerve cord (VA10, VB11, DB7, AS10 and DA7). Wild type GFP::FUST-1 localized predominantly to the nucleus and had an overall nuclear index of 0.9 ± 0.02. GFP::FUST-1 R524S and P525L proteins had overall nuclear indices of 0.21 ± 0.02 and - 0.12 ± 0.02, respectively. Data collected from three independent trials. Error bars indicate -± SEM. Neurons of the same genotype were pooled for pairwise comparison. Pairwise t-test: * *p* < 0.05; ** *p* < 0.01; *** *p* < 0.001.

Transgenes expressing *C. elegans* FUST-1 protein with an N-terminal GFP fusion tag (GFP::FUST-1) were used to determine the nuclear-to-cytoplasmic ratio of GFP::FUST-1 in *C. elegans* motor neurons (Figure 1B). For this analysis, animals were mechanically immobilized with microbeads and GFP::FUST-1 fluorescence was measured in five cholinergic motor neurons (VA10, VB11, DB7, AS10 and DA7). As expected [24], wild type *C. elegans* GFP::FUST-1 localized predominantly to the nucleus, while GFP::FUST-1 R524S and P525L proteins showed cytoplasmic displacement (Figure 1C). We calculated the nuclear index of GFP fluorescence for each reporter (Figure 1D). The predominantly nuclear GFP::FUST-1WT protein had a nuclear index of 0.9 ± 0.02 per neuron (Figure 1E). By contrast, GFP::FUST-1 R524S and P525L proteins had nuclear indices of 0.21 ± 0.02 and −0.12 ± 0.02, respectively, consistent with cytoplasmic displacement (Figure 1E, **Figures S2 A and B**). We conclude that the nuclear localization of *C. elegans* FUST-1 protein in motor neurons is disturbed when ALS-causal mutations are inserted, replicating localization changes seen in FUS ALS patient motor neurons.

In patient motor neurons, FUS mutations lead to cytoplasmic FUS deposits [2]. By contrast, live transgenic *C. elegans* immobilized with microbeads showed diffuse cytoplasmic GFP::FUST-1R24S and GFP::FUST-1P525L fluorescence in motor neurons. However, when animals were briefly immobilized with 2,3-butanedione monoxime (BDM) or sodium azide (NaN3), GFP::FUST-1 formed inclusions in motor neurons (**Figure S2 C**). These chemicals are frequently used to anesthetize *C. elegans*, but they induce metabolic [25] or mitochondrial stress [26], respectively. Thus, stress may drive formation of cytoplasmic GFP-FUST-1 inclusions. To confirm, we treated animals with dithiothreitol (DTT) to induce endoplasmic reticulum (ER) stress [27]. Brief DTT exposure induced cytoplasmic FUST-1 inclusions in motor neurons expressing the R524S and P525L GFP:FUST-1 proteins, but not in animals overexpressing wild type GFP::FUST-1 (**Figure S2 D and E**). Consistent with observations in other systems, our results suggest that stress drives formation of FUS inclusions [9,24].

### Generation of *C. elegans* knock-in ALS *fust-1* models

To generate *C. elegans* ALS FUS knock-in models, we used template-driven, CRISPR/Cas9-mediated genome editing to insert R524S and P525L equivalent mutations into the C-terminal NLS of the endogenous *fust-1* gene on chromosome II (Figure 2A). Because genome editing requires introduction of silent codon changes, we generated a wild type control strain containing silent mutations (Figure 2B). A “C” superscript (*fust-1WT^C^*, *fust-1R524S^C^*, and *fust-1P525L^C^)* is used to distinguish the edited alleles from the control, non-transgenic *fust-1(+)* allele found in the standard laboratory *C. elegans* strain (N2). We refer to *fust-1R524S^C^* and *fust-1P525L^C^* animals together as ALS *fust-1* animals.

**Figure 2.**
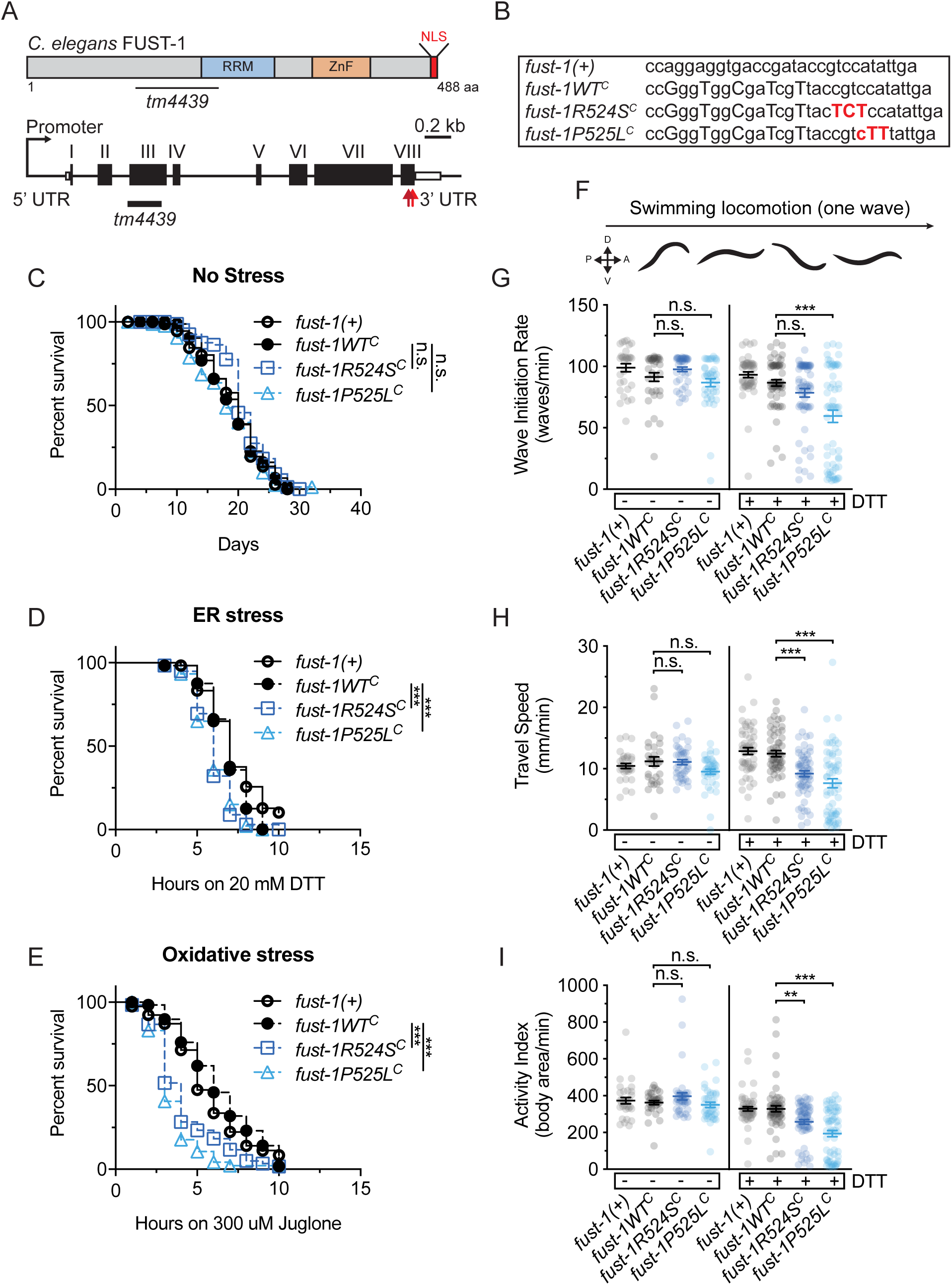
Generation and characterization of *C. elegans* ALS FUS knock-in models. **(A)** Illustration of *C. elegans* FUST-1 protein (top) and ALS *fust-1* knock-in models (bottom) generated by CRISPR/Cas9-mediated homologous recombination. The endogenous *fust-1* locus on chromosome II was edited using CRISPR/Cas9-mediated homologous recombination to knock in nuclear localization signal mutations corresponding to R524S and P525L (red arrows). A control wild type *fust-1WT^C^* was generated in tandem containing all the silent codon changes necessary for CRISPR/Cas9 genome editing. All alleles generated by CRISPR/Cas9 are annotated with a “C” superscript. RRM: RNA recognition motif. ZnF: Zinc Finger domain. NLS: Nuclear Localization Signal. **(B)** Mutations induced by CRISPR/Cas9-mediated homologous recombination in ALS *fust-1* and control animals. Capitalized letters show all the mutations (both silent and missense) inserted by genome editing. Red bases show the location of ALS associated missense mutations. **(C-E)** Under standard culture conditions, ALS *fust-1* animals had normal lifespan. ALS *fust-1* animals were treated with 20 mM DTT or 300 uM juglone to induce ER or oxidative stress, respectively. ALS *fust-1* knock-in alleles decreased survival on both types of stress compared to *fust-1WT^C^* controls. Three independent trials. Error bars indicate -SEM. Log-rank test: *p* < 0.001.-- **(F-I)** ALS *fust-1* animals have latent locomotion defects after exposure to DTT-induced ER stress. Swimming locomotion was assessed with computer vision software CeleST [29]. ALS *fust-1* animals reared under standard culture conditions have normal locomotion. ALS *fust-1* alleles lead to locomotion defects after a brief period of DTT-induced ER stress compared to *fust-1WT^C^* animals. Data collected from three independent trials. Error bars indicate SEM. Pairwise t-test: ** *p* <0.01; *** *p* < 0.001.

FUS overexpression leads to severe locomotion defects [9,12,13,28]. ALS *fust-1* knock-in animals had no overt locomotion defects on solid surfaces, or detectable swimming defects on day 1 or on day 10 of adulthood (Figure 2F-I and **Figure S3 A**, using CeleST [29]). In addition, *C. elegans* FUS knock-in models had normal life spans (Figure 2C), pharyngeal feeding (**Figure S3 B**), and no neuron loss was observed.

### Knock-in animals are hypersensitive to stress and have stress-induced locomotion defects

We have previously shown that stress reveals latent defects in *C. elegans* knock-in models for ALS SOD1 [30]. ALS FUS mutations may also impair cellular stress responses [24,31,32][30]. To identify latent defects in ALS FUS animals, we examined the impact of DTT-induced ER stress [27] and juglone-induced oxidative stress [33]. ALS *fust-1* animals had decreased survival after stress, compared to *fust-1WT^C^* controls. To examine if loss of *fust-1* function contributes to stress sensitivity, we examined stress response in *fust-1(tm4439)* loss of function animals [34], referred to as *fust-1(-)* animals. Loss of *fust-1* sensitized animals to DTT, but not to juglone (**Figure S3 C**). Thus, loss of *fust-1* function may contribute to the ER stress sensitivity, but not oxidative stress sensitivity, in ALS *fust-1* knock-in models.

To reveal latent stress-induced defects, we tested swimming locomotion after DTT-induced ER stress (Figure 2F-I). Swimming ALS *fust-1* animals had impaired locomotion compared to *fust-1WT^C^* animals after a brief ER stress exposure (Figure 2F-I, CeleST [29]). Combined, our analysis of ALS FUS animals shows that stress induces formation of FUST-1 inclusions within *C. elegans* motor neurons, and reveals latent survival and locomotion defects.

### Knock-in animals show impaired autophagy in motor neurons and muscles, m

Recent evidence suggests that wild type FUS overexpression may impair neuronal autophagy [11]. However, it remains unclear whether ALS FUS mutations impair neuronal autophagy. To monitor autophagy, we used a tandem-tagged dual GFP::mCherry::LGG-1 reporter broadly expressed under the *lgg-1* promoter [35]. LGG-1 is the *C. elegans* ortholog of mammalian Atg8, a component of autophagy membranes. Accumulation of the GFP::mCherry::LGG-1 reporter into autophagy membranes leads to formation of green+red fluorescent autophagosomes (APs) (Figure 3A). Fusion of green+red fluorescent APs with acidic lysosomes quenches pH-sensitive GFP fluorescence, resulting in red fluorescent autolysosomes (ALs) (Figure 3A). Finally, lysosomal enzymes degrade the GFP::mCherry::LGG-1 protein, abolishing all fluorescence.

**Figure 3.**
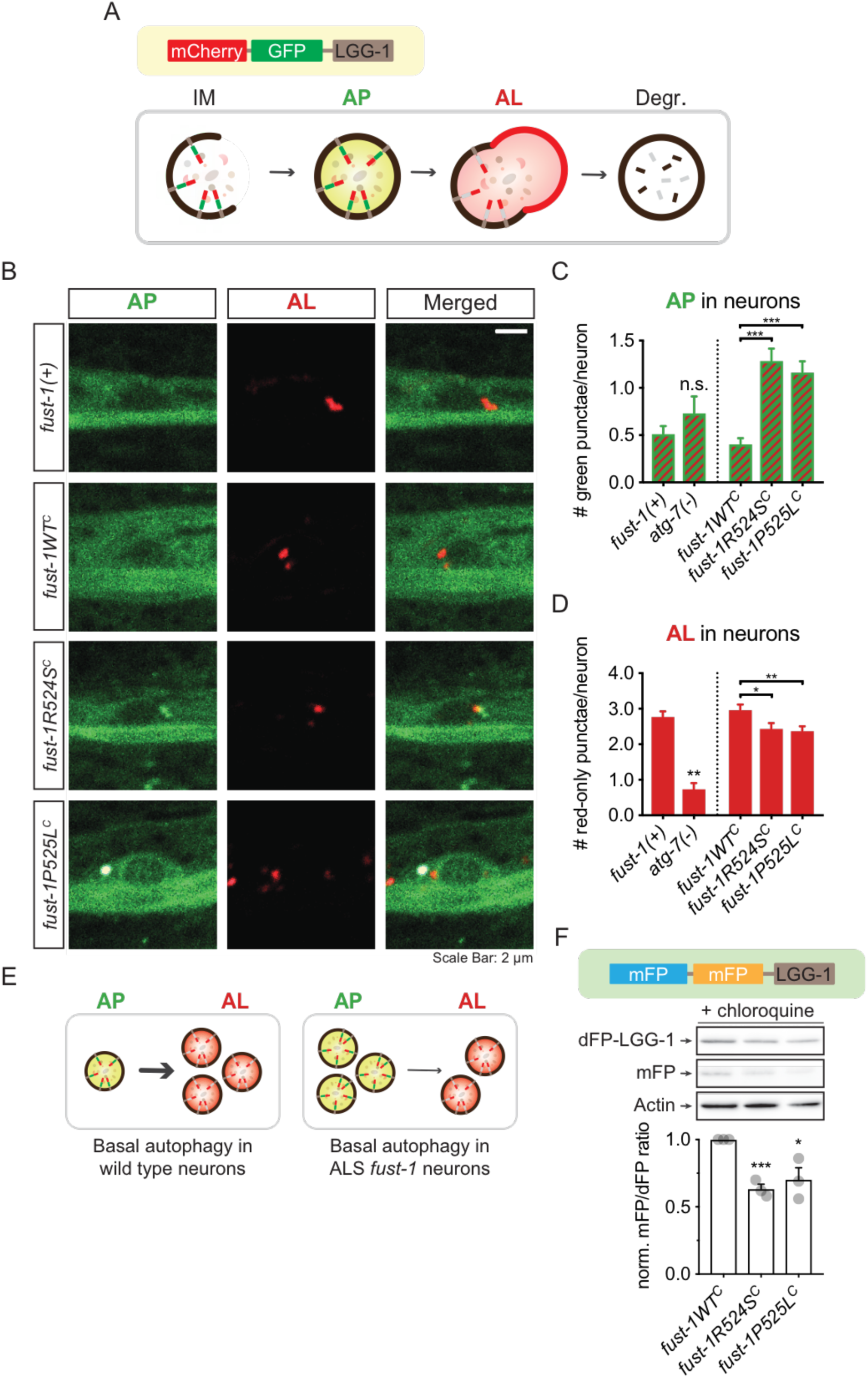
Analysis of autophagy in motor neurons of ALS *fust-1* animals. **(A)** Illustration: Animals were mechanically paralyzed to assess mCherry::GFP::LGG-1 reporter accumulation on isolation membranes (IM) and autophagosomes (AP). Green+Red dual-labelled autophagosome (AP) fusion with acidic lysosomes quenches GFP; autolysosomes (AL) have only red fluorescence. Fluorescence disappears after degradation (Degr.) in the lysosome. **(B)** Representative images of day 1 old adult ALS *fust-1* animals and controls expressing the mCherry::GFP::LGG-1 reporter in motor neurons. Scale Bar = 2 um. **(C)** APs and ALs in the motor neurons of ALS *fust-1* animals and controls. *atg-7(bp411)* loss of function animals are defective in IM maturation [35], but this had no impact on the number of green+red AP punctae *versus* non-transgenic *fust-1(+)* N2 animals. ALS *fust-1* alleles had increased green+red AP punctae compared to *fust-1WT^C^* controls. **(D)** Loss of *atg-7* function decreased red AL punctae *versus* non-transgenic *fust-1(+)* N2 animals. ALS *fust-1* alleles decreased the number of red AL punctae in motor neurons compared to *fust-1WT^C^* controls. In C and D, error bars indicate -SEM. Pairwise t-test: * *p* < 0.05; ** *p* < 0.01; *** *p* < 0.001.-- **(E)** Schematic representation: basal neuronal autophagy in ALS *fust-1* animals is perturbed with improper accumulation of APs and reduction in the number of ALs. **(F)** Western Blot using a different neuronally expressed LGG-1 transgene double-tagged with two mono-fluorescent (mFP) proteins: dFP::LGG-1 (top). Lysosomal hydrolases cleave the sequence between mFPs. Ratio of mFP to dFP::LGG-1 protein levels measures autophagy [36]. Middle: representative Westerns for dFP::LGG-1, mFP and actin from ALS *fust-1* and *fust-1WT^C^* control animals, after overnight chloroquine treatment. Chloroquine inhibits lysosomal activity [36], allowing for visualization of neuronal mFP proteins in autolysosomes. Without chloroquine treatment, the mFP protein band was invisible. Bottom: neuronal mFP/dFP::LGG-1 protein ratio normalized against *fust-1WT^C^* controls. ALS *fust-1* alleles decrease mFP/dFP::LGG-1 ratio compared to *fust-1WT^C^* controls. Three independent trials. Error bars indicate -± SEM. Pairwise t-test: * *p* < 0.05; *** *p* < 0.001. Results in 3C and D were collected at the same time as the data presented in Figure 5D and E, and are from the same *fust-1(+)*, *fust-1WT^C^*, *fust-1R524S^C^* and *fust-1P525L^C^* animals.

As a control, we first examined neuronal autophagy in the motor neurons of *atg-7(bp411)* loss of function animals (Figure 3B and 3C). *atg-7* function is required early in autophagy [35]. As predicted, loss of *atg-7* function diminished the number of neuronal ALs compared to non-transgenic N2 animals (Figure 3C) [35]. We found that ALS *fust-1* alleles increased APs and decreased ALs in motor neurons, compared to *fust-1WT^C^* controls. We confirmed neuronal autophagy defects in ALS *fust-1* animals by rearing animals under different conditions and saw qualitatively similar autophagy defects (**Figure S4 A-D**). Combined, these results suggest that neuronal autophagy activity is impaired in ALS *fust-1* animals (Figure 3D).

Neuronal autophagy defects in ALS *fust-1* animals were also quantified using a different LGG-1 transgene double-tagged with two monomeric fluorescent proteins (Cerulean and Venus) expressed under the neuronal *rab-3* promoter [36]. The double fluorescent LGG-1 protein, referred to as dFP::LGG-1, localizes to autophagic membranes. Degradation of dFP::LGG-1 in lysosomes generates transient monomeric fluorescent proteins (mFP). Comparison of relative mFP to dFP::LGG-1 protein levels by Western Blot permits the quantitation of neuronal autophagy flux [36]. We found that ALS *fust-1* alleles decreased the relative ratio of neuronal mFP to full-length dFP::LGG-1, suggesting that neuronal autophagy flux is impaired in ALS *fust-1* animals (Figure 3F). Taken together, our findings suggest that ALS *fust-1* mutations disrupt basal neuronal autophagy.

Next, we determined if loss of *fust-1* function causes similar defects in autophagy. Visual inspection of GFP::mCherry::LGG-1 fluorescence in motor neurons (**Figure S5 A-C**), or quantification of neuronal mFP to dFP::LGG-1 protein levels by Western Blot (**Figure S5 D**), did not reveal autophagy defects in *fust-1(-)* animals. Combined, our data suggest that ALS *fust-1* alleles cause a gain of toxic function that perturbs neuronal autophagy.

We also examined if autophagy defects in ALS FUS knock-in models are specific to neurons. Visual inspection of GFP::mCherry::LGG-1 fluorescence in body wall muscles revealed similarly increased levels of APs compared to *fust-1WT^C^* controls (**Figure S6 A-C**). Consequently, ALS *fust-1* alleles perturb both neuronal and muscular autophagy.

ALS *fust-1* alleles could globally disrupt all major cellular catabolic pathways, including the ubiquitin proteasome system (UPS). However, we did not detect UPS defects in two different experimental paradigms (**Figure S7**). Thus, ALS *fust-1* alleles specifically impair autophagy, while leaving the UPS intact.

### Knock-in animals have impaired neuromuscular function

Neuromuscular dysfunction precedes motor neuron degeneration in ALS patient motor neurons [37–39]. To test for neuromuscular defects in ALS *fust-1* knock-in models, we examined resistance to aldicarb, a pharmacological inhibitor of acetylcholinesterase [40]. Aldicarb exposure causes paralysis due to excessive cholinergic excitation at the neuromuscular junction (NMJ) and eventually leads to death (Figure 4A) [40]. Decreased synaptic release delays paralysis onset, while increased release can accelerate paralysis [40]. Notably, overexpression of ALS-linked human FUS delta57 in *C. elegans* GABAergic neurons causes aldicarb hypersensitivity [10]. We found that ALS FUS knock-in animals were also hypersensitive to aldicarb-induced paralysis and immobilized more quickly than *fust-1WT^C^* animals in the presence of 1mM aldicarb (Figure 4B). No changes in NMJ protein localization were observed (**Figure S8 A-D**). *fust-1(-)* animals were not markedly sensitive to aldicarb, although a trend toward hypersensitivity was noted (Figure 4B). We conclude that ALS FUS animals show impaired neuromuscular function.

**Figure 4.**
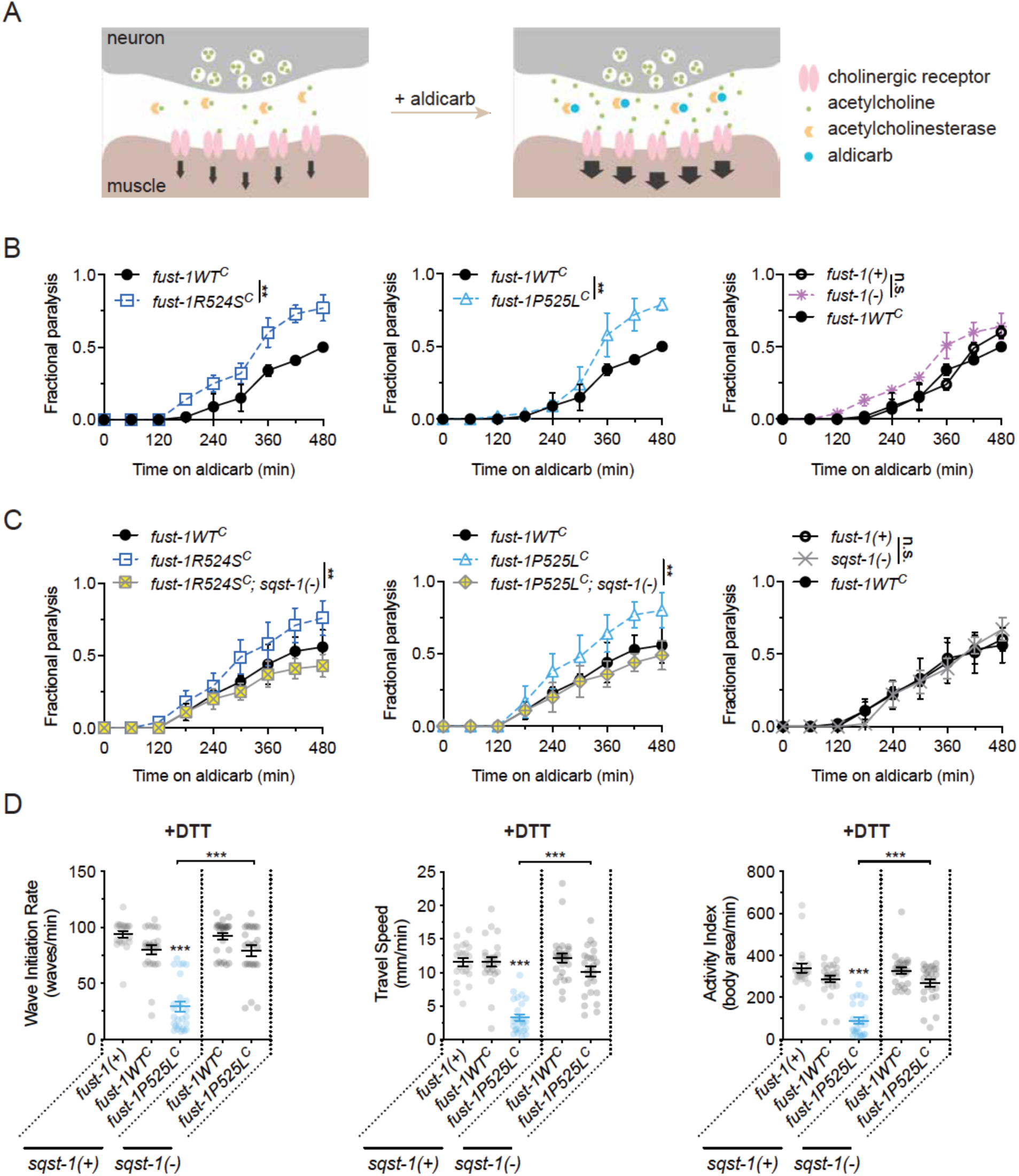
ALS *fust-1* alleles disrupt neuromuscular function. Loss of *sqst-1*, *C. elegans* ortholog for mammalian SQSTM1, suppresses neuromuscular function defects and stress-induced locomotion defects in ALS *fust-1* animals. **(A)** Neuromuscular function was assessed with aldicarb, an acetylcholinesterase inhibitor. Inhibition of acetylcholinesterase leads to acetylcholine buildup at the neuromuscular junction, resulting in hyperexcitation of the postsynaptic muscle and paralysis. While wild type animals on aldicarb paralyze with a characteristic time-course; hypersensitivity or resistance to aldicarb suggests impaired neuromuscular function. **(B)** ALS *fust-1* animals are hypersensitive to aldicarb and paralyze faster compared to *fust-1WT^C^* animals. Loss of *fust-1* function did not significantly change aldicarb sensitivity in *fust-1(tm4439)* loss of function animals. The data for all three graphs was collected at the same time, and *fust-1WT^C^* in each graph represents the same data. The data is presented in different graphs for clarity. Three independent trials. Error bars indicate SEM. Log-rank test: * *p* < 0.05; ** *p* <0.01; *** *p* < 0.001. **(C)** Loss of *sqst-1*, the *C. elegans* ortholog for SQSTM1, suppresses aldicarb hypersensitivity in ALS *fust-1* animals. The data for all three graphs was collected at the same time, and *fust-1WT^C^* in the first two graphs represents the same data. The data is presented in different graphs for clarity. Three independent trials. Error bars indicate SEM. Log-rank test: * *p* < 0.05; ** *p* <0.01; *** *p* < 0.001. **(D)** Loss of *sqst-1* function partially suppresses DTT-induced locomotion defects in *fust-1P525L^C^* animals. Swimming locomotion was assessed with computer vision software CeleST. Data was collected from two independent trials. Error bars indicate SEM. Pairwise t-test: * *p* < 0.05; ** *p* <0.01; *** *p* < 0.001.

Increased aldicarb sensitivity could also arise from alterations in muscular function, which we examined using levamisole resistance assay. Exposure to levamisole, a cholinergic receptor agonist, leads to paralysis due to excessive body wall muscle contraction (**Figure S9 A**). Mutations that impair muscular function or structure can alter responses to levamisole. ALS *fust-1* animals were hypersensitive to levamisole compared to *fust-1WT^C^* animals, suggesting muscular function is also impaired in ALS FUS models (**Figure S9 B**).

### Loss of *C. elegans* SQSTM1 suppresses locomotion and neuromuscular function defects in knock-in animals

SQSTM1/p62 selectively recognizes and delivers cargo to autophagy membranes for degradation [41]. SQSTM1 accumulates in ALS FUS patient motor neurons and SQSTM1 mutations can cause ALS. We examined the impact of loss of *sqst-1*, the *C. elegans* SQSTM1 ortholog, on neuromuscular and locomotion defects in ALS *fust-1* animals using the *sqst-1(ok2892)* loss of function allele. First, we examined aldicarb sensitivity in double-mutant animals. Loss of *sqst-1* function rescued aldicarb hypersensitivity in both *fust-1R524S^C^* and *fust-1P525L^C^* animals (Figure 4C). However, *sqst-1* loss of function by itself had no impact on aldicarb resistance, suggesting *sqst-1* function may be dispensable for neuromuscular function in wild type animals. Another *sqst-1* loss of function allele partially rescued aldicarb hypersensitivity in *fust-1P525L^C^* animals (**Figure S10 A**; *sqst-1(ok2869)*).

We also examined stress-induced locomotion defects in double mutant animals. After DTT-induced ER stress, the locomotion defects in *fust-1P525L^C^* animals were partially suppressed by *sqst-1* loss of function. (Figure 4C). However, loss of *sqst-1* function did not rescue levamisole hypersensitivity in ALS *fust-1* animals (**Figure S9 C**). Consequently, ALS *fust-1* alleles may impair neuronal and muscular function via independent mechanisms.

### Loss of *C. elegans* SQSTM1 does not restore mutant FUST-1 localization

Loss of *C. elegans* SQSTM1 ortholog might alleviate ALS *fust-1* neuromuscular function defects by restoring nuclear *fust-1* localization. We tested this hypothesis using transgenic animals expressing wild type or mutant *C. elegans* FUST-1 protein with an N-terminal GFP fusion tag (GFP::FUST-1) in *sqst-1(ok2892)* loss of function background. Loss of *sqst-1* function did not restore mutant GFP::FUST-1 predominant nuclear localization in the motor neurons (Figure 5A-B). And, loss of *sqst-1* function did not alter nuclear localization of wild type FUST-1 protein (Figure 5A-B). Thus, loss of *sqst-1* function does not suppress aldicarb response defects by correcting cytoplasmic mislocalization of the mutant FUST-1 protein in *C. elegans* motor neurons.

**Figure 5.**
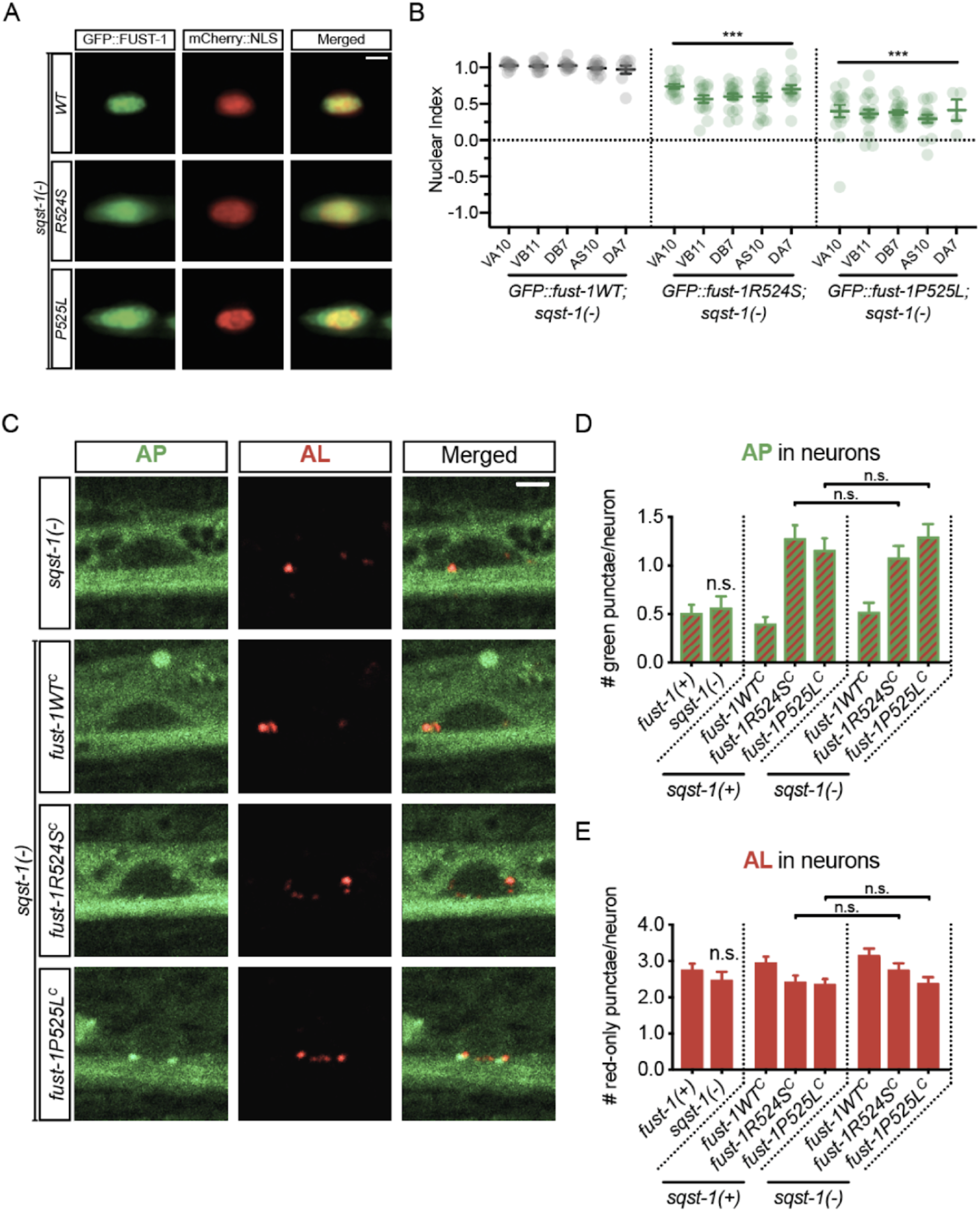
Loss of SQST-1 function does not restore nuclear FUST-1 protein localization or suppress autophagy dysfunction in ALS FUS models. **(A)** Representative images of wild type or mutant GFP::FUST-1 proteins in motor neurons of *sqst-1* loss of function animals. Transgenes driving the expression of wild type or mutant *C. elegans* FUST-1 protein with an N-terminal GFP fusion tag (GFP::FUST-1) were used to determine relative nuclear-to-cytoplasmic GFP::FUST-1 localization in the motor neurons of *sqst-1* loss of function animals. The neuronal nuclei were labelled with mCherry::NLS driven under the *cho-1* cholinergic promoter. Images of mechanically-immobilized animals were taken at a focal plane where the neuronal nuclei could be clearly observed. Loss of SQST-1 function does not restore nuclear localization of either mutant GFP::FUST-1 protein. Scale bar = 2 um. **(B)** Nuclear index of wild type or mutant GFP::FUST-1 proteins in motor neurons of *sqst-1* loss of function animals was calculated as shown in Figure 1. **(C)** Representative images of day 1 adult *sqst-1(ok2892)* loss of function animals and ALS *fust-1*, *sqst-1* loss of function double mutant animals expressing the mCherry::GFP::LGG-1 reporter in motor neurons. Scale bar = 2 um. **(D)** Quantification of autophagosomes (AP) in the motor neurons of *sqst-1* loss of function and ALS *fust-1* double mutant animals, and controls. Error bars indicate -SEM. Pairwise t-test. **(E)** Quantification of ALs in the motor neurons of *sqst-1* loss of function and ALS *fust-1* double mutant animals, and controls. Error bars indicate -SEM. Pairwise t-test. Results in D and E were collected at the same time as the data presented in Figure 3C and D, and are from the same *fust-1(+)*, *fust-1WT^C^*, *fust-1R524S^C^* and *fust-1P525L^C^* animals.

### Loss of *C. elegans* SQSTM1 does not rescue neuronal autophagy flux

Alternatively, loss of the *C. elegans* SQSTM1 ortholog might alleviate ALS *fust-1* neuromuscular function defects by restoring autophagy function. Thus, we wondered if ALS *fust-1* alleles perturb neuronal autophagy in a *sqst-1* dependent manner. To test this hypothesis, we examined APs and ALs labelled by the GFP::mCherry::LGG-1 reporter in double-mutant animals carrying an ALS *fust-1* mutation and the *sqst-1(ok2892)* loss of function allele. However, loss of *sqst-1* function did not suppress ALS *fust-1* autophagy defects in motor neurons (Figure 5C-E). Furthermore, loss of *sqst-1* function alone in otherwise wild type animals had no impact on the APs or ALs in motor neurons (Figure 5C-E). These results suggest that SQST-1 is dispensable for autophagy in *C. elegans* motor neurons. We conclude that ALS *fust-1* alleles do not disrupt neuronal autophagy by perturbing SQST-1 function.

### Exposure to aldicarb induces stress in *C. elegans*

The responses of ALS *fust-1* animals to aldicarb suggests that this treatment might be stressful to *C. elegans*. To assess this, we examined the subcellular localization of DAF-16 FOXO transcription factor in wild type animals treated with aldicarb. Cellular stress leads to nuclear translocation DAF-16, driving transcription of stress response genes [42]. Aldicarb exposure increased the nuclear translocalization of DAF-16::GFP in the intestines of wild type animals compared to control animals, suggesting that aldicarb treatment is stressful for *C. elegans* (Figure 6A). Indeed, aldicarb treatment for four hours led to formation of GFP::FUST-1 neuronal inclusions (**Figure S11 A and B**). We suggest that aldicarb exposure is stressful and may drive neuronal stress in ALS *fust-1* animals.

**Figure 6.**
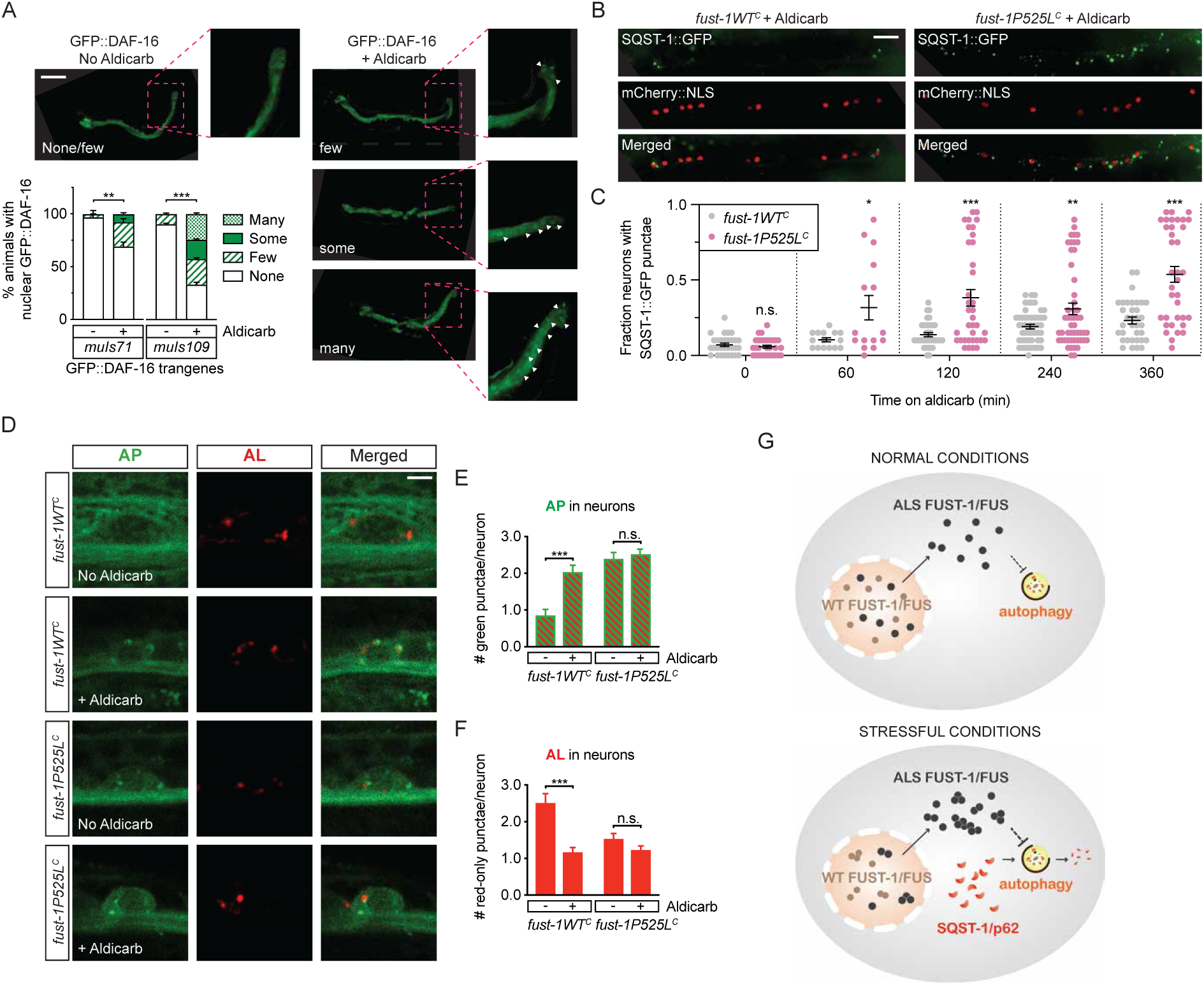
Aldicarb treatment causes SQST-1 accumulation in motor neurons of ALS FUS animals. **(A)** Aldicarb treatment induces nuclear translocation of GFP::DAF-16. Images: representative of none, few, some or many intestinal cells showing predominant nuclear GFP::DAF-16 localization before and after 4 hours of 1 mM aldicarb treatment. Animals were categorized according to nuclear GFP::DAF-16 localization levels, ranging from none to many. White arrows indicate nuclear-localized GFP::DAF-16 in posterior intestinal cells of *muIs109*; *daf-16(mu86)* animals (expanded images, pink lines). Scale bar = 150 um. Graph: Quantification of GFP::DAF-16 nuclear localization in *daf-16(mu86)* loss of function animals with two different GFP::DAF-16 transgenes: *muIs71* and *muIs109*. Two independent trials. Chi-square test: ** *p* < 0.01; *** *p* < 0.001. **(B)** Aldicarb exposure and SQST-1::GFP accumulation in motor neurons. Without aldicarb, both *fust-1P525L^C^* and *fust-1WT^C^* animals show low levels of SQST-1::GFP accumulation (not shown). Left image: after 4 hours of 1mM aldicarb treatment, *fust-1WT^C^* control animals accumulated little *fust-1WT^C^*. Right image: But, in *fust-1P525L^C^* animals the fraction of motor neurons with accumulation sometimes dramatically increased. Little SQST-1::GFP accumulation was seen in muscles, regardless of aldicarb treatment. Cholinergic motor neurons are labelled with nuclear-localized mCherry. SQST-1::GFP was expressed under the *sqst-1* promoter. Scale bar = 10 um. **(C)** SQST-1::GFP accumulation in motor neurons after aldicarb treatment. Each dot represents the fraction of motor neurons with SQST-1::GFP accumulation for one animal. Aldicarb treatment increased SQST-1::GFP accumulation in *fust-1P525L^C^* animals *versus fust-1WT^C^*. At least two independent trials per time point. N = 15 animals per genotype at 1 hour. N > 30 animals per genotype for the remaining time points. Error bars indicate SEM. Pairwise t-test: * *p* < 0.05; ** *p* <0.01; *** *p* < 0.001. **(D)** Aldicarb exposure changes neuronal autophagy. Representative images of neuronal autophagy in day 1 adult *fust-1WT^C^* and *fust-1P525L^C^* animals expressing the mCherry::GFP::LGG-1 reporter. Scale bar = 2 um. **(E and F)** Aldicarb-induced autophagy changes in motor neurons of day 1 adult *fust-1WT^C^* and *fust-1P525L^C^* animals. Four hours of aldicarb exposure increased autophagosome (AP) levels and decreased autolysosome levels (AL) in control *fust-1WT^C^* animals. Without aldicarb, *fust-1P525L^C^* animals showed increased AP levels and decreased AL levels (also Figure 3B-D). After aldicarb exposure, AP and AL levels did not substantively change. N > 46 animals per genotype. Pairwise t-test: * *p* < 0.05; ** *p* < 0.01; *** *p* < 0.001. **(G)** Summary. Top: mutant FUST-1 diffusely mislocalizes into the cytoplasm under standard culture conditions, and a gain of toxic function mechanism disrupts basal neuronal autophagy. Bottom: Under stressful conditions, mutant FUST-1 forms cytoplasmic inclusions and induces improper SQST-1 accumulation.

### SQST-1 accumulates specifically in motor neurons of ALS *fust-1* animals after perturbations of neuromuscular function by aldicarb

SQSTM1/SQST-1 protein accumulates with age and stress [43]. Furthermore, FUS and SQSTM1 proteins co-aggregate in the motor neurons of ALS FUS patients, suggesting that mutant FUS protein leads to accumulation and/or aggregation of SQSTM1 in neurons. To test this hypothesis, we crossed a SQST-1::GFP reporter into *fust-1WT^C^* and *fust-1P525L^C^* knock-in animals and examined SQST-1::GFP accumulation in cholinergic motor neurons. The SQST-1::GFP levels were relatively low in the motor neurons of animals reared under normal culture conditions, and the fraction of motor neurons with SQST-1::GFP accumulation was similarly low at 0.07 ± 0.011 and 0.058 ± 0.01 in *fust-1WT^C^* and *fust-1P525L^C^* animals, respectively (Figure 6B and 6C). Interestingly, aldicarb treatment over time induced higher levels of SQST-1::GFP accumulation in the motor neurons of *fust-1P525L*^C^ animals compared to *fust-1WT^C^* animals (Figure 6B and 6C). Levamisole treatment did not induce neuronal SQST-1::GFP accumulation in either *fust-1P525L*^C^ animals or *fust-1WT^C^* controls (**Figure S11 C**). SQSTM1 is degraded through autophagy and improper SQSTM1 accumulation may indicate autophagy defects [42]. Since loss of *sqst-1* function suppresses neuromuscular defects, but not autophagy defects, we conclude that neuronal SQST-1 accumulation likely acts downstream of autophagy dysfunction in ALS *fust-1* animals.

### Exposure to aldicarb changes neuronal autophagy flux in *C. elegans*

The results above suggest that aldicarb treatment may exacerbate neuronal autophagy defects in ALS FUS animals. This drove us to ask if aldicarb treatment induces changes in neuronal autophagy of normal animals. We examined the GFP::mCherry::LGG-1 reporter in motor neurons after 4 hours of aldicarb treatment. Aldicarb treatment of *fust-1WT^C^* animals elevated the steady state levels of APs and decreased ALs levels (Figure 6D-F). By contrast, aldicarb treatment did not exacerbate apparent neuronal autophagy defects in *fust-1P525L^C^* animals compared to untreated *fust-1P525L^C^* controls (Figure 6D-F), possibly due to a ceiling effect associated with the preexisting autophagy defects in *fust-1P525L^C^* animals. Thus, aldicarb treatment elicits stress responses and alters autophagy activity in *C. elegans* motor neurons.

Combined, our results suggest that cytoplasmic ALS *fust-1* alleles impair basal autophagy in *C. elegans* motor neurons (Figure 6G). Since we did not observe the same autophagy defects in *fust-1* loss of function animals, these defects are likely driven by a gain of toxic function in cytoplasmically mislocalized ALS FUST-1 protein. Furthermore, aldicarb-induced perturbations of neuromuscular function elicit a stress response, change neuronal autophagy, and cause disproportionate SQST-1 accumulation in ALS *fust-1* animal motor neurons (Figure 6G). Thus, stress reveals latent autophagy defects in the motor neurons of ALS *fust-1* animals.

## Discussion

Together, the ubiquitin proteasome and the autophagy-lysosome systems help maintain proteostasis by degrading proteins. Accumulation of ubiquitin and SQSTM1 positive inclusions in ALS FUS patient motor neurons suggests defective proteolysis contributes to pathogenesis [44]. However, early processes that lead to aberrant protein localization and aggregation in ALS FUS patient motor neurons remain elusive. Here, we show that ALS FUS knock-in alleles in *C. elegans* perturb basal neuronal autophagy, while leaving the ubiquitin proteasome system (UPS) intact. Furthermore, loss of function in SQSTM1 ortholog *sqst-1*, an ALS-associated autophagy adaptor protein, suppressed neuromuscular function defects and stress-induced locomotion defects in ALS FUS knock-in models. Our findings suggest a role for autophagy defects in ALS FUS-mediated neuronal dysfunction.

Here, we found that ALS FUS knock-in mutations perturbed basal autophagy in *C. elegans* motor neurons. Visual inspection of GFP::mCherry::LGG-1 reporter in ALS FUS model motor neurons showed induction of autophagosomes (APs) with a concurrent depletion in autolysosomes (ALs). Similarly, lysosomal degradation of a neuronal dFP::LGG-1 reporter was diminished in ALS FUS models. We conclude that neuronal autophagy is perturbed in ALS FUS knock-in animals. Consistent with this notion, a recent study found elevated numbers of LC3-positive autophagosomes in ALS FUS patient motor neurons, independently corroborating autophagy dysfunction in ALS FUS pathogenesis [23]. Neuronal autophagy defects in ALS FUS knock-in animals do not fully mimic neuronal autophagy defects in early autophagy mutants suggesting that ALS *fust-1* alleles may perturb different and/or additional steps in the autophagy pathway. While more research is necessary to better understand autophagy in motor neurons and the precise step(s) at which neuronal autophagy is perturbed in ALS FUS models, we suggest that autophagy dysfunction may contribute to early decline in protein homeostasis in ALS FUS pathogenesis.

ALS FUS knock-in alleles may have cell-type specific impacts on autophagy. As seen in *C. elegans* motor neurons, *C. elegans* muscles had increased APs in ALS FUS knock-in animals. However, the number of ALs in muscles was not changed. Differences in neuronal and muscular autophagy in ALS FUS models may arise in part from tissue-specific variations in basal autophagy. Indeed, two previous studies showed marked differences in *C. elegans* neuronal and muscular autophagy [35,36].

Disease-associated mutations in FUST-1 lead to cytoplasmic FUST-1 mislocalization in *C. elegans* motor neurons [10,14]. Therefore, ALS FUS knock-in alleles may perturb neuronal autophagy through cytoplasmic gain of function and/or nuclear loss of function in *fust-1*. However, we found that neuronal autophagy in *fust-1* loss of function animals was largely intact. Combined, our findings suggest that a gain of toxic function causes defects in neuronal autophagy. Consistent with this, overexpression of wild type FUS protein in neuronal cell lines alternately increases and decreases autophagosome and autolysosome levels, respectively [11]. In addition, transgenic mice overexpressing wild type FUS show SQSTM1 accumulation in motor neurons, suggesting autophagy dysfunction [11]. Combined, our results complement previous work in FUS overexpression models and suggest that a gain of toxic function in mutant FUS disrupts neuronal protein homeostasis.

ALS patient motor neurons show cytoplasmic FUS inclusions by disease end stage [21]. However, in *C. elegans* motor neurons, mutant FUST-1 proteins were diffusely mislocalized into the cytoplasm and did not form inclusions under standard culture conditions. Lack of cytoplasmic FUST-1 inclusions may speak to the relative young age of adult *C. elegans* compared to human patients, who accumulate cytoplasmic FUS inclusions in motor neurons after decades [21]. Still, our findings are consistent with other work that shows i) human ALS FUS protein overexpression in *C. elegans* neurons does not lead to inclusion formation in young adult animals [9] and ii) FUS inclusions are dispensable for neuronal dysfunction in an ALS-linked FUS knock-in model in mice [14]. Consequently, FUS mislocalization may be an early event in disease pathogenesis. We note that examined strains herein were not fixed prior to visualization, thus allowing for the observation of neuronal autophagy reporters in live, unstressed animals.

Synaptic dysfunction and denervation preceed neuron loss in ALS patients [38,39,45]. Here, we assessed neuromuscular function in ALS FUS knock-in animals by examining response to aldicarb, an acetylcholinesterase inhibitor. ALS *fust-1* animals, unlike *fust-1* loss of function animals, were hypersensitive to aldicarb, consistent with a gain of toxic function in ALS *fust-1* perturbing NMJ function. Notably, overexpression of ALS-linked human FUS delta57 in *C. elegans* GABAergic neurons also leads to aldicarb hypersensitivity [10]. While our results support a gain of toxic function in ALS FUS, animals lacking *fust-1* also showed modest, albeit not significant, hypersensitivity to aldicarb. Therefore, we hesitate to rule out a role for *fust-1* loss of function in the neuromuscular dysfunction of ALS FUS knock-in models.

ALS-linked FUS differentially localizes to cytoplasmic stress granules after exposure to stress [46,47] and may perturb cellular stress responses. Here, we found that ER stress led to formation of cytoplasmic FUST-1 inclusions in transgenic animals expressing mutant, but not wild type, FUST-1, suggesting that stress may reveal latent defects in ALS FUS knock-in animals. Consistent with this, ER stress and oxidative stress decreased survival in *C. elegans* ALS FUS knock-in models and ER stress led to locomotion defects. ALS FUS knock-in animals are hypersensitive to aldicarb and aldicarb treatment induced nuclear translocation of FOXO transcription factor DAF-16 in the intestines of wild type animals.Therefore, we suggest that aldicarb treatment induces an organismal level stress response. Combined, our findings suggest that ALS FUS knock-in mutations sensitize animals to various stressors.

Our findings also suggest that neuronal autophagy changes as part of normal physiological responses to stress in wild type animals. Aldicarb modulates neuronal autophagy in wild type animals by increasing and decreasing AP and AL levels, respectively. It is unclear if these steady-state changes represent increased or decreased neuronal autophagy activity. However, induction of APs and depletion of ALs suggests impediment of neuronal autophagy flux after aldicarb treatment. By comparison, aldicarb treatment failed to induce a similar response in ALS *fust-1* knock-in animals. This may be due to a ceiling effect; ALS *fust-1*-induced neuronal autophagy defects are present even without aldicarb treatment. Aldicarb treatment also lead to inappropriate accumulation of SQST-1 in ALS *fust-1* motor neurons. Since both SQSTM1 and SQSTM1-bound cargo are degraded primarily through autophagy, aldicarb treatment likely induces disproportionate SQST-1 accumulation in ALS *fust-1* motor neurons due to enhanced defects in neuronal autophagy. We conclude that stress likely exacerbates ALS FUS-induced defects in protein homeostasis.

In patient motor neurons, FUS aggregates with SQSTM1. Furthermore, mutations in SQSTM1 have been linked to ALS [21]. Thus, FUS and SQSTM1 may share disease mechanisms [44]. Here, we also examined the impact of loss of function in SQSTM1 ortholog *sqst-1* in ALS FUS knock-in models. Remarkably, loss of *sqst-1* suppressed neuromuscular and ER stress-induced locomotion defects in ALS FUS knock-in models. Taken together, our findings suggest that ALS *fust-1* alleles perturb neuronal autophagy independent of SQST-1 function. While we do not yet know how SQSTM1/SQST-1 accumulation perturbs neuronal function, stress and SQSTM1/SQST-1 accumulation in autophagy-deficient ALS FUS knock-in animals likely further compromise proteostasis networks. Taken together, our findings in *C. elegans* ALS FUS knock-in models show that FUS knock-in alleles perturb neuronal autophagy and disrupt protein homeostasis. Furthermore, aldicarb-induced changes in neuromuscular function induce accumulation of SQSTM1 ortholog in *C. elegans*. And, while loss of *C. elegans* SQSTM1 suppresses neuromuscular defects in ALS FUS knock-in models, our results demonstrate that autophagy defects are upstream of, and not dependent on, SQSTM1 dysfunctions. Future work is needed to define how ALS FUS-induced changes in neuronal autophagy may compromise cellular stress response pathways, and ultimately perturb neuronal function.

## Materials and Methods

### Cloning

Cloning for wild type or mutant GFP::FUST-1 overexpression in cholinergic neurons: For *Punc-17::GFP::fust-1::unc-54 3’ UTR* plasmids, four PCR products with 20-30 bp overlap (a 4506 bp cholinergic *unc-17* promoter, a 906 bp GFP fragment, a 2619 bp *fust-1* DNA (wild type or mutant) and 388 bp *unc-54 3’ UTR* fragment) were assembled into KpnI and SpeI double-digested pBlueScript using NEBuilder High DNA Assembly Cloning Kit (E5520S). Equivalent ALS FUS patient mutations for R524S and P525L are at the end of the last *fust-1* exon, and thus fall within the overlapping sequence between the last two assembled fragments (*fust-1* DNA fragment and *unc-54 3’ UTR* fragment). Consequently, different primers were used to PCR amplify and assemble *fust-1* + *unc-54 3’ UTR* fragments in wild type *versus* mutant *Punc-17::GFP::fust-1::unc-54 3’ UTR* plasmids. The *unc-17* promoter, GFP and *unc-54* 3’ UTR fragments were PCR amplified from pHA#757 (*Punc-17::GFP::unc-54 3’UTR*) plasmid [48] with primer pairs unc-17p_f and unc-17p_r, GFP_f and GFP_r, and an allele specific *unc-54 3’ UTR* forward primer unc-54_3UTR_fust-1_f and unc-54_3UTR_fust-1_r, respectively. The *fust-1* DNA fragments (wild type or mutant) were amplified from ALS *fust-1* knock-in model genomic DNA using fust-1_ex1_f and an allele specific *fust-1* exon 8 fust-1_ex8_r reverse primers. In addition, an in-frame 27 bp flexible linker sequence (GGAGCATCGGGAGCCTCAGGAGCATCG) was inserted between GFP and *fust-1* DNA fragments, by including parts of it in the GFP_r and fust-1_ex1_f primers. These primers are listed in **Table S1**. The resulting plasmids pHA#820 (*Punc-17::GFP::linker::fust-1WT::unc-54 3’ UTR)*, pHA#821 (*Punc-17::GFP::linker::fust-1R524S::unc-54 3’ UTR), and pHA#822 (Punc-17::GFP::linker::fust-1P525L::unc-54 3’ UTR*) were verified by sequencing.

Cloning for CRISPR/Cas9-mediated *fust-1* knock-in: A U6 promoter-driven *fust-1* guide RNA was amplified from the *PU6::klp-12::sgRNA* (Addgene plasmid #46170) vector template by replacing the *klp-12* targeting sequence with *fust-1* specific primer pair *fust-1_guide_f* and *fust-1_guide_r*. The resulting DNA was circularized to generate pHA#823 *pU6::fust-1::guide*.

### Strain generation and maintenance

Strains were maintained at 20°C under standard culture conditions, unless otherwise indicated. To generate animals expressing GFP::FUST-1 in cholinergic motor neurons, *Punc-17::GFP::fust-1::unc-54 3’ UTR* plasmids carrying wild type or mutant *fust-1* allele were injected at 25 ng/ul into non-transgenic N2 animals with 100 ng/ul of pBlueScript vector and 2.5 ng/ul of PCFJ90 *Pmyo-2::mCherry* (Addgene plasmid #19327). The *Pmyo-2::mCherry* co-injection marker was used to maintain extrachromosomal arrays driving GFP::FUST-1 expression in motor neurons.

ALS *fust-1* alleles *R524S^C^* and *P525L^C^* were introduced into the endogenous *fust-1* locus on chromosome II using CRISPR/Cas9-mediated homologous recombination and *pha-1(ts)* co-conversion as in [49]. The guide RNA construct pHA#823 *pU6::fust-1::guide* was injected at 50 ng/ul into temperature-sensitive GE24 (*pha-1(e2123) III*) animals with 50 ng/ul of *Peft-3::Cas9*, 50ng/ul of pJW1285 *pha-*1 sgRNA, 10 uM of 200mer sense *pha-1(+)* rescue oligo [49] and 10uM of a mutation specific single-stranded oligodeoxynucleotide listed in **Table S1**. Silent codon changes were introduced to generate a PvuI restriction site, and to prevent the re-hybridization of guide RNAs with edited *fust-1* alleles. A wild type *fust-1WT^C^* control containing silent mutations only was also generated. Putative recombinants among *pha-1(+)* rescued animals were screened for insertion events with PCR amplification using fust-1_scr_f and and fust-1_scr_r, followed by PvuI digestion. Isolated alleles were verified by sequencing and backcrossed four times. The endogenous *pha-1* locus was PCR amplified and sequenced to verify the replacement of repaired *pha-1* allele with wild type *pha-1(+)* in all strains.

### Behavioral and survival assays

#### Survival

Animals reared under standard culture conditions at 20 °C and scored on alternating days starting from the first day of adulthood. FUDR was not used [50]. Aging animals were transferred to a new seeded plate every other day until all animals stopped laying eggs. Animals unresponsive to touch were scored as dead. To score survival on dithiothreitol (DTT), 20 mM DTT (Thermo Fisher Scientific R086) plates were prepared the day before the assay. To score survival on 5-Hydroxy-1,4-naphthoquinone (juglone), fresh 300 uM juglone (Sigma Aldrich H47003) plates were prepared on the same day as the survival assay. In both DTT and juglone sensitivity assays, animals were scored every hour for survival until all animals became unresponsive to touch.

#### Pharyngeal feeding/pumping

Day 1 adults were transferred onto NGM plates seeded with 200 ul of fresh bacterial OP50 lawn and feeding was recorded for 10 seconds per animal. Videos were scored manually.

#### Swimming

Animals on day 1 and day 10 of adulthood were transferred into 60 ul of M9 pipetted onto a 1 mm diameter ring on a microscope slide. Swimming was recorded at 18 frames per second for 30 seconds. Videos were analyzed using CeleST [29].

#### Aldicarb and levamisole resistance

Day 1 adults were transferred onto 1 mM aldicarb (Sigma-Aldrich 33386) or 0.4 mM Levamisole (Sigma L9756) NGM plates. Paralysis was scored every hour for 8 hours by sequential prodding twice to the tail and twice to the head with a platinum pick. Animals were scored as paralyzed if no movement or pharyngeal pumping was observed for 5 seconds after prodding. Paralyzed animals were censored and were not scored again for subsequent time points. Fractional paralysis over time was calculated and data was analyzed with GraphPad Prism (La Jolla, CA) using log-rank analysis to establish statistical significance.

### Neuronal survival

Motor neuron survival was examined in transgenic animals on day 1 of adulthood using a cholinergic (*cho-1::mCherry*) neuronal marker. A full list of strains used in this study can be found in **Table S2**. Day 1 adult animals were mounted on 15% (vol/vol) agarose (IBI Scientific) pads and immobilized with 0.1 micron Polybead Polystyrene Microspheres (Polysciences #9003-53-6). Fluorescent neurons were visualized and scored at the microscope for cell death based on loss of neuronal mCherry under 63x or 100x objectives (Zeiss AxioImager ApoTome and AxioVision software v4.8).

Glutamatergic neuron degeneration was examined by retrograde dye-filling with DiD (Fisher DilC18(5) D307) for 1.5 hours. Animals were spun down at 10000 rpm for 1 min, and transferred to a regular NGM plate. After 1 hour, animals were mounted on 15% (vol/vol) low melting point agarose pads (IBI Scientific) pads and immobilized with 0.1 micron Polybead Polystyrene Microspheres (Polysciences #9003-53-6). Fluorescent neuronal cell bodies were visualized and scored for lack of dye uptake under 63x or 100x objectives (Zeiss AxioImager ApoTome and AxioVision software v4.8).

### Synaptic puncta imaging

Day 1 adult animals were mounted on 2% (vol/vol) agar pads and paralyzed using 30 mg/mL 2-3-butanedione monoxime (BDM) (Sigma) in M9 buffer. Images were captured in z-stacks from dorsal cord posterior to vulva with a 100x objective, using Zeiss Axio Imager ApoTome Microscope and AxioVision software v4.8. For synaptic puncta imaging, we used BDM rather than microbeads to immobilize animals, as mechanically immobilized animals contract their muscles during imaging, which changes synaptic puncta locations in consecutive z-stack images. Data from at least three independent trials (*n* ≥ 19 animals in total/genotype for all groups) was analyzed. Puncta total intensity and linear density were quantified using the Punctaanalyser program in Matlab (v6.5; Mathworks, Inc., Natick, MA, USA; RRID:SCR_001622).

### Quantification of subcellular GFP::FUST-1 localization and inclusions

Animals expressing a wild type or mutant *C. elegans* FUST-1 protein with an N-terminal GFP fusion tag (GFP::FUST-1) were used to determine the relative nuclear-to-cytoplasmic GFP::FUST-1 localization in cholinergic motor neurons. *Pcho-1::mCherry::NLS* was used to visualize cholinergic motor neurons. For quantitative analysis, animals were mounted with microspheres as described above. GFP::FUST-1 fluorescence was measured in five motor neurons along the ventral nerve cord (VA10, VB11, DB7, AS10 and DA7) under a 100x objective (Zeiss AxioImager ApoTome and AxioVision software v4.8). Images were taken at a focal plane where the neuronal nuclei could be clearly observed. Relative nuclear and cytoplasmic fluorescence intensities were determined with ImageJ. Nuclear index of GFP::FUST-1 was calculated by first taking the difference between nuclear and cytoplasmic GFP fluorescence in a single neuron, and dividing by total GFP fluorescence in the same neuron. The average and SEM were calculated and data was analyzed with GraphPad Prism (La Jolla, CA). For quantification of GFP::FUST-1 accumulations in neurons, animals were paralyzed in the same way, and neuronal cytoplasmic and nuclear punctae were manually scored in the same five motor neurons along the ventral nerve cord, after 15 minutes of 20 mM DTT treatment or after 4 hours of 1 mM aldicarb treatment.

### Quantification of SQST-1 accumulations by microscopy

A *sqst-1::GFP* fusion reporter driven under the *C. elegans sqst-1* promoter was used to visualize SQST-1 accumulation in motor neurons. Cholinergic motor neurons along the ventral nerve cord were visualized with a cholinergic *Pcho-1::mCherry::NLS* neuronal marker. To determine the number of motor neurons with SQST-1 accumulations, we scored 20 cholinergic motor neurons posterior to vulva in day 1 adult animals mechanically immobilized on microspheres as described above. SQSTM1 accumulations were scored under normal culture conditions, after 1 mM aldicarb treatment for one, two, four or six hours, and after 0.4 mM levamisole treatment for 4 hours. Motor neurons with SQSTM1 accumulations were manually scored at the microscope under a 40x objective (Zeiss AxioImager ApoTome and AxioVision software v4.8). The average and SEM were calculated and data was analyzed with GraphPad Prism (La Jolla, CA).

### Autophagy by microscopy

To visualize autophagy activity in motor neurons, we used animals expressing the dual-tagged GFP::mCherry::LGG-1 reporter. Day 1 adults were mounted live with microspheres. The images were acquired with Olympus FV3000 Confocal Laser Scanning inverted microscope. GFP excitation and emission were set to 488 nm and 500-540 nm, respectively. mCherry excitation and emission were set to 561 nm and 570-670 nm, respectively. mCherry fluorescence intensity is greater and the gain of red channel was set lower to avoid overexposure, as previously described by Chang *et al.* [35]. Consequently, the cytoplasmic GFP::mCherry::LGG-1 reporter appears green in the images. Ventral cord motor neurons were identified based on location and morphology. Z-stack images of the motor neurons were acquired at 60x using 0.34 um steps over a vertical distance of 9.52 um. Green+red and red-only punctae were manually counted in the ventral nerve cord motor neurons in one 0.34 um slice where the neuronal nuclei could be observed. To visualize autophagic activity in body wall muscles, transgenic animals expressing the dual-tagged GFP::mCherry::LGG-1 reporter were similarly immobilized on day 1 of adulthood. Z-stack images of body wall muscles were acquired at 60x using 0.34 um steps over a vertical distance of 9.52 um. Green+red and red-only punctae were manually counted in body wall muscles in one 0.34 um slice immediately below the slice with muscle filaments. The average and SEM were calculated, and data was analyzed with GraphPad Prism (La Jolla, CA).

In an independent set of experiments, we confirmed neuronal autophagy defects. Animals were raised on bacteria expressing double-stranded RNA against GFP and *squat-1* (a.k.a. *sqt-1*) on NGM plates prepared with 100 ug/ml ampicillin and 6 mM isopropyl β-D-1-thiogalactopyranoside (IPTG). L4 stage animals were passaged onto RNAi plates seeded with 200 ul of 10X concentrated bacterial strains L4417 and B0491.2 [51] to drive IPTG-induced RNAi against GFP and *sqt-1*, respectively. While feeding RNAi induces downregulation of respective genes in non-neuronal tissues, *C. elegans* neurons are resistant to feeding RNAi [52]. Thus, feeding RNAi disproportionately increased the GFP::mCherry::LGG-1 reporter signal-to-noise ratio in motor neurons (**Figure S4**). Feeding RNAi against *sqt-1* partially suppressed the rolling phenotype in dual-tagged GFP::mCherry::LGG-1 reporter lines, facilitating inspection of motor neurons After four days on RNAi plates, L4 stage progeny were transferred onto fresh RNAi plates. The next day, animals on day 1 of adulthood were tested for autophagy activity in neurons. All bacterial clones were verified by sequencing.

### Neuronal autophagy by Western Blot

To assess neuronal autophagy flux by Western Blot, we integrated a tandem-tagged neuronal LGG-1 dual fluorescent reporter (strain DLM11 from [36]) and crossed it into ALS *fust-1* models. To synchronize different genetic populations, 100 gravid adults were allowed to lay eggs for 5 hours at 20°C on large 60 ml NGM plates seeded with 2.4 ml of OP50. At the end of 5 hours, gravid adults were removed from the plates and progeny were grown at 20°C for 55 hours. L4 stage progeny were then washed off the plates with 10 ml of M9, placed in 15 ml tubes, and incubated with 4x concentrated OP50 in 20 mM chloroquine diphosphate salt (Sigma-Aldrich) for 18 hours. Animals were then washed six times with 10 ml of M9. To collect the samples, animals were spun down at 4000 rpm for 1 min and resuspended in 50 ul of M9. Samples were then added with 150 ul of RIPA buffer (150 mM NaCl, 0.1 % SDS, 0.5 % sodium deoxycholate, 1 % Triton and 1M Tris-HCl, pH 8) containing a protease inhibitor cocktail (Sigma Aldrich Complete™, Mini, EDTA-free Protease Inhibitor Cocktail Tablet), frozen in liquid nitrogen, and crushed with a mortar and pestle to break up the cuticle. Finally, samples were transferred into microcentrifuge tubes, further homogenized by sonication at 30 Hz and spun at 14,000 rpm at 4ºC for 15 minutes to pellet out the debris. Protein concentration in the supernatant was determined with Pierce BCA Protein Assay Kit (Thermo Scientific 23225). Protein lysates were heated at 95ºC for 10 minutes and separated by 12% sodium dodecyl sulfate–polyacrylamide gel electrophoresis (SDS-PAGE). The proteins were then transferred onto polyvinylidene difluoride (PVDF) membranes with a 20% Methanol transfer buffer solution for 1.5 hours. Membranes were blocked for 1 hour in 5% milk dissolved in Tris-buffered saline with Tween-20 (TBST, containing 10 mM Tris, 150 mM NaCl, and 0.1% Tween-20, pH 7.6), followed by incubation with primary antibodies 1:1000 mouse anti-GFP (Sigma-Aldrich 11814460001) and 1:5000 mouse anti-Actin (Abnova MAB8172) in 5% bovine serum dissolved in TBST at 4ºC overnight. Membranes were washed with TBST three times for 15 minutes per wash and incubated with 1:5000 HRP-conjugated rabbit anti-mouse secondary antibody (Abcam ab6728) in 5% milk+0.5% bovine serum albumin dissolved in TBST for 1 hour. Membranes were again washed x 3 with TBST for 15 minutes per wash, then visualized with SuperSignal West Pico Chemiluminescent Substrate (Thermo Scientific 34077) using a C600 Azure Biosystems imaging system (Dublin, CA). Densitometry analysis was conducted with ImageJ software.

### Ubiquitin Proteasome System (UPS) activity assay

Day 1 adults were mounted live on 15% (vol/vol) low melting agarose (IBI Scientific) pads in M9 medium and immobilized with 1 mM levamisole (Sigma L9756) dissolved in M9. Images were acquired with Olympus FV3000 Confocal Laser Scanning inverted microscope with sequential line scanning. Images were acquired periodically every 5 minutes over one hour following photoconversion of a single neuron. Images of individual neurons were acquired with a 60x 1.3 numerical aperture (NA) objective and Olympus silicone immersion oil. Photoconversion experiments on individual neurons were carried out with a 405 nm 50 mW laser at 1% laser output with a scanning speed of 200 us per pixel. We used a 488 nm, 20 mW and 561 nm, 20 mW lasers to visualize UbG76V::Dendra2. The relative green and red UbG76V::Dendra2 fluorescence intensities over time were normalized against the green and red UbG76V::Dendra2 fluorescence intensity observed immediately before and after photoconversion, respectively. We manually adjusted the focus in-between time points as necessary. Fluorescence intensities were analyzed on ImageJ.

## Author Contributions

S.N.B.

N.C.

B.U.

J.L.

K.S.

J.S. A.C.H.

## Acknowledgements

We thank L. Lapierre for providing us with *atg-7(sp411); lgg-1::GFP::mCherry*. Funding from ALS Finding A Cure, ALS Association. Kelsey Schuch was supported by OVPR Brown University Seed Award research funds. Burak Unsal was supported by Kirac Foundation. Natalie Chapkis and Jonah Simon were supported by Brown University Karen T. Romer Undergraduate Teaching and Research Awards.

