## Supplemental Material for "Disrupted Autophagy and Neuronal Dysfunction in *C. elegans* Knock-in Models of FUS Amyotrophic Lateral Sclerosis"

### Supplemental Figures and Tables

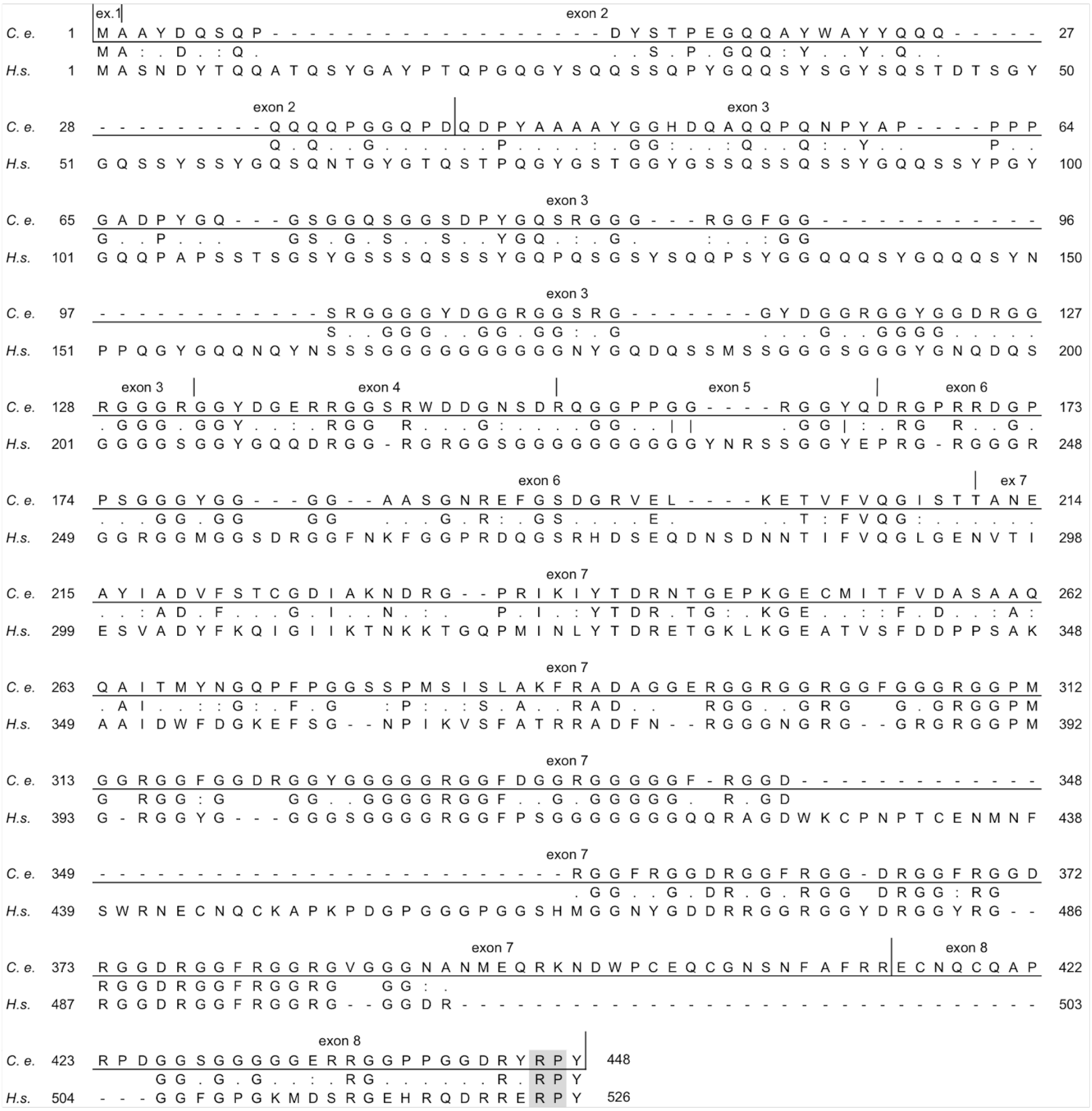


**Figure S1. Alignment of complete human FUS and *C. elegans* FUST-1 protein sequences (related to Figure 1).** Identical amino acid residues are indicated between protein sequences; conserved and semi-conserved amino acid residues are indicated with ":" and ".", respectively. Majority of ALS-linked mutations disrupt the C' terminal PY nuclear localization signal (PY-NLS, aa 507-526), encoded by the last exon of FUS. *C. e.* for *C. elegans* FUST-1 protein GenPept Accession Number NP_495483.1; *H.s.* for *Homo sapiens* FUS GenPept Accession Number CAG33028.1. Exon boundaries are from *C. elegans fust-1* transcript C27H5.3a.1. ex: exon.

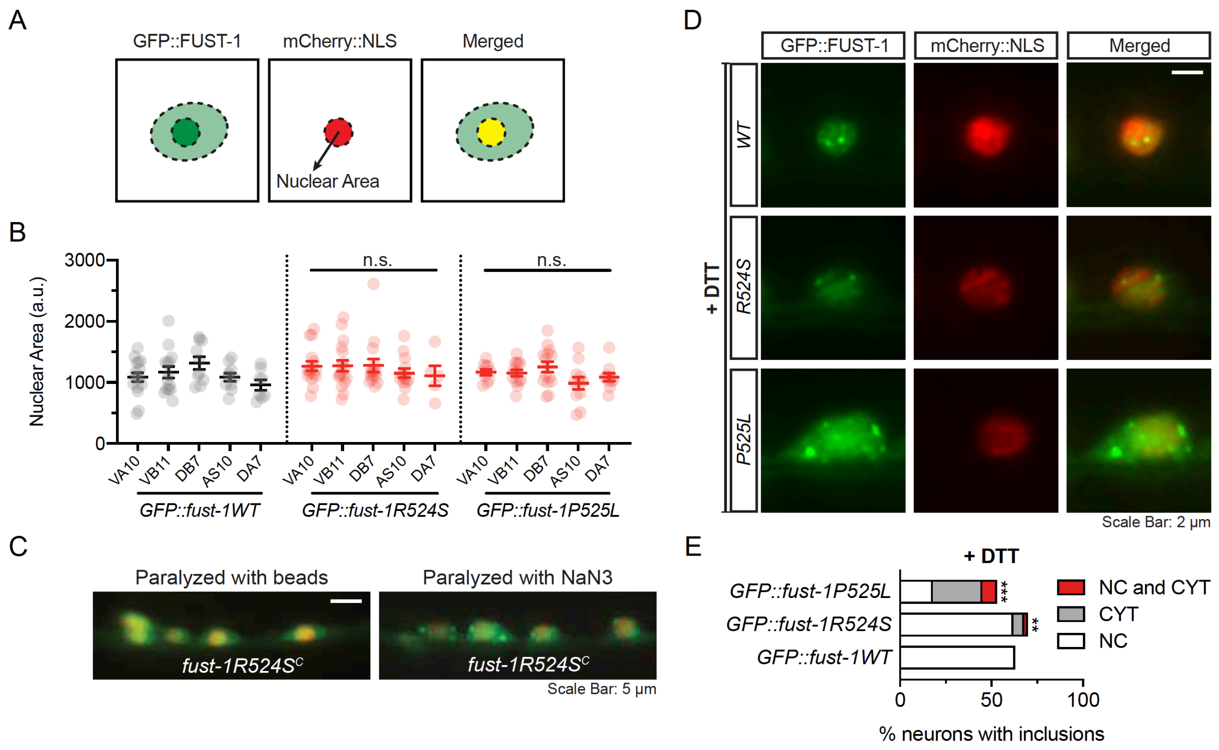


**Figure S2. Stress induces cytoplasmic FUST-1 inclusion formation (related to Figure 1).**

**(A-B)** Quantification of the nuclear area labelled by the mCherry::NLS reporter in motor neurons expressing wild type or mutant GFP::FUST-1. No change in nuclear area or morphology was observed in motor neurons expressing the wild type or mutant GFP::FUST-1 protein. Data collected from three independent trials. N ≥ 58 neurons per genotype. Error bars indicate -± SEM. Neurons of the same genotype were pooled for pairwise comparison. Pairwise t-test.

**(C)** Representative images of transgenic animals expressing mutant GFP::FUST-1 in motor neurons after they have been immobilized with microbeads or sodium azide (NaN3). Pharmacologically anesthetized animals show nuclear and cytoplasmic mutant GFP::FUST-1 inclusions in motor neurons. Similarly, transgenic animals expressing wild type GFP::FUST-1 in motor neurons showed nuclear-localized inclusions after animals were paralyzed with Sodium Azide (NaN3) (not shown).

**(D-E)** Transgenic animals expressing wild type GFP::FUST-1 in motor neurons showed nuclear-localized FUST-1 inclusions after Dithiothreitol (DTT)-induced ER stress. The same treatment induced formation of both nuclear and cytoplasmic inclusions in motor neurons of animals expressing mutant GFP::FUST-1. Three independent trials. N > 19 for each genotype. NC: nuclear. CYT: cytoplasmic. Chi-square test: ** $p < 0.01$; *** $p < 0.001$.

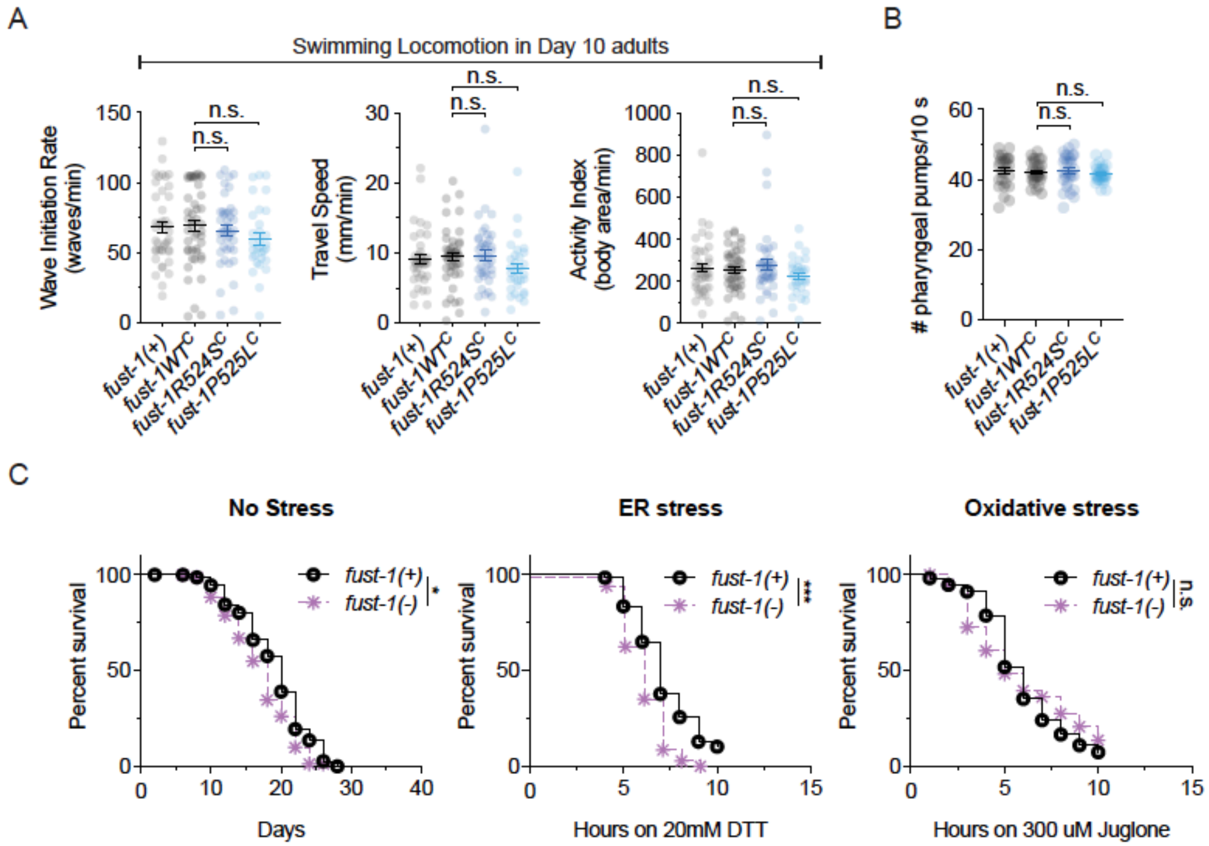


**Figure S3. ALS FUS knock-in animals have normal swimming locomotion on day 10 of adulthood and have normal pharyngeal feeding. Loss of endogenous *fust-1* function decreases lifespan, and sensitizes animals to ER stress (related to Figure 2).**

**(A)** ALS *fust-1* animals on day 10 of adulthood have normal locomotion. Swimming locomotion was assessed with computer vision software CeleST [29]. N > 30 animals per genotype. Data collected from three independent trials. Error bars indicate -SEM. Pairwise t-test.

**(B)** ALS *fust-1* animals on day 1 of adulthood have normal pharyngeal pumping. N > 27 animals per genotype. Data collected from three independent trials. Error bars indicate -SEM. Pairwise t-test.

**(C)** *fust-1(tm4439)* loss of function allele sensitizes animals to DTT-induced ER stress, but not to juglone-induced oxidative stress. N = 30 animals per genotype in each trial. Three independent trials. Error bars indicate -SEM. Log-rank test: * $p < 0.05$; ** $p < 0.01$; *** $p < 0.001$.--

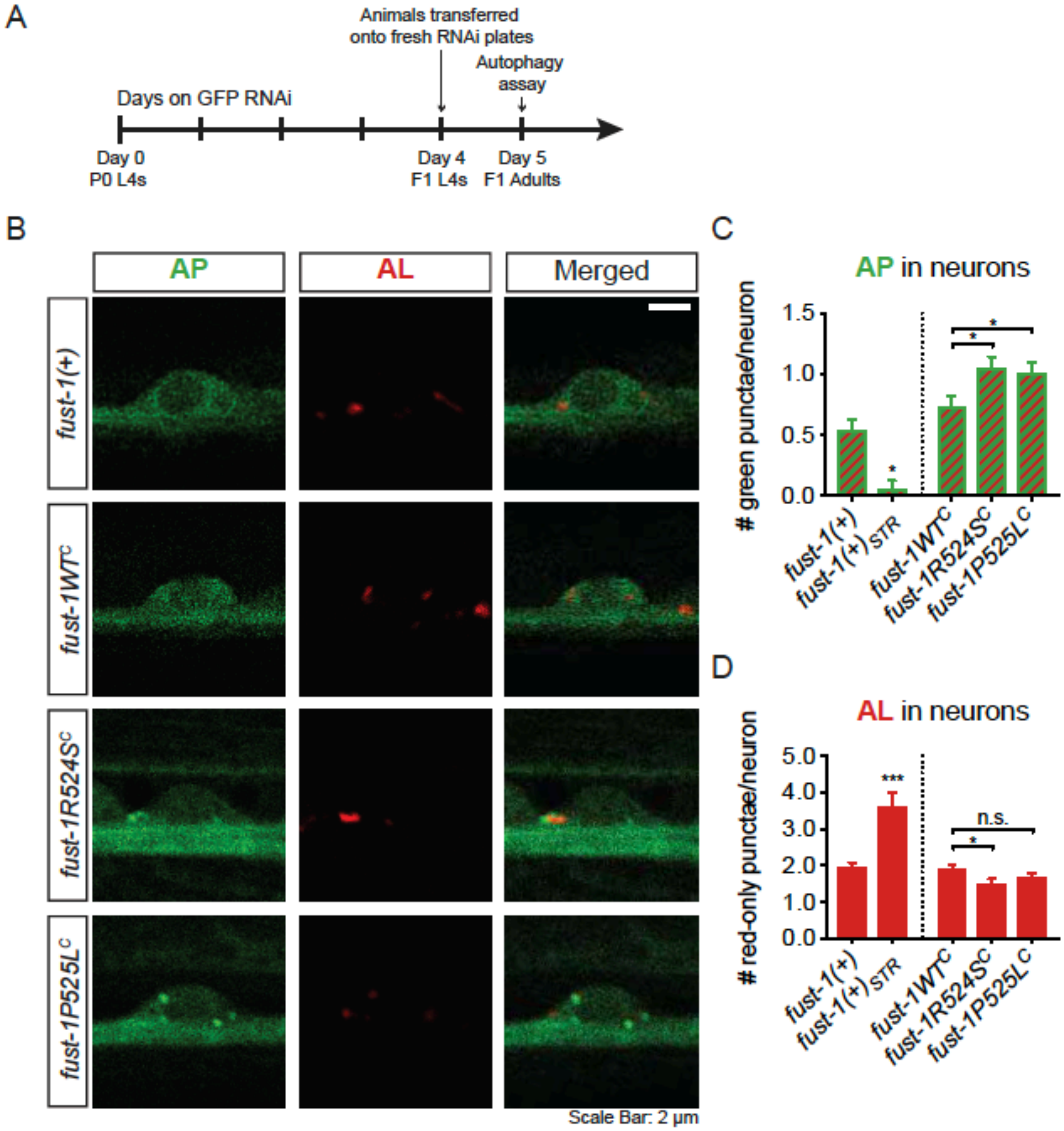


**Figure S4. Examination of mCherry::GFP::LGG-1 autophagy reporter in motor neurons of ALS *fust-1* animals raised on HT115 RNAi bacteria (related to Figure 3).**

**(A)** To decrease mCherry::GFP::LGG-1 signal in non-neuronal tissues and straighten animal posture, transgenic animals were grown on RNAi bacterial strains transcribing double-stranded RNA (dsRNA) against GFP and *squat-1* mRNA. The nutritional composition of RNAi bacteria differs from OP50 [53]. L4 stage progeny were transferred onto fresh RNAi plates the day before the assay. The next day, animals on day 1 of adulthood were examined for neuronal autophagy. *squat-1* RNAi partially suppressed the rolling phenotype in mCherry::GFP::LGG-1 transgenic animals, allowing for the visualization of motor neurons.

**(B)** Representative images of ALS *fust-1* animals and controls on day 1 of adulthood expressing mCherry::GFP::LGG-1 reporter in motor neurons following feeding RNAi against GFP and *sqt-1*. Scale bar = 2 um.

**(C)** Quantification of green+red APs in the motor neurons of ALS *fust-1* animals and controls after feeding RNAi. As a control, we examined neuronal autophagy in starved animals. Starvation diminished the number of green+red AP punctae in motor neurons compared to satiated non-transgenic *fust-1(+)* N2 animals. Contrarily, ALS *fust-1* alleles increased the number of APs per neuron compared to *fust-1WT*$^{C}$ controls. N > 116 neurons per genotype. Data collected from five independent trials. Error bars indicate SEM. Pairwise t-test: * $p < 0.05$; ** $p < 0.01$; *** $p < 0.001$. *fust-1(+)*$_{STR}$: overnight starved animals.

**(D)** Quantification of red-only ALs in the motor neurons of ALS *fust-1* animals and controls after feeding RNAi. As a control, we examined neuronal autophagy in starved animals. Starvation elevated the number of red-only AL punctae in motor neurons compared to satiated non-transgenic *fust-1(+)* animals. Contrarily, ALS *fust-1R524S*$^{C}$ alleles decreased the number ALs per neuron compared to *fust-1WT*$^{C}$ controls. N > 116 neurons per genotype. Data collected from five independent trials. Error bars indicate SEM. Pairwise t-test: * $p < 0.05$; ** $p < 0.01$; *** $p < 0.001$. *fust-1(+)*$_{STR}$: overnight starved animals.

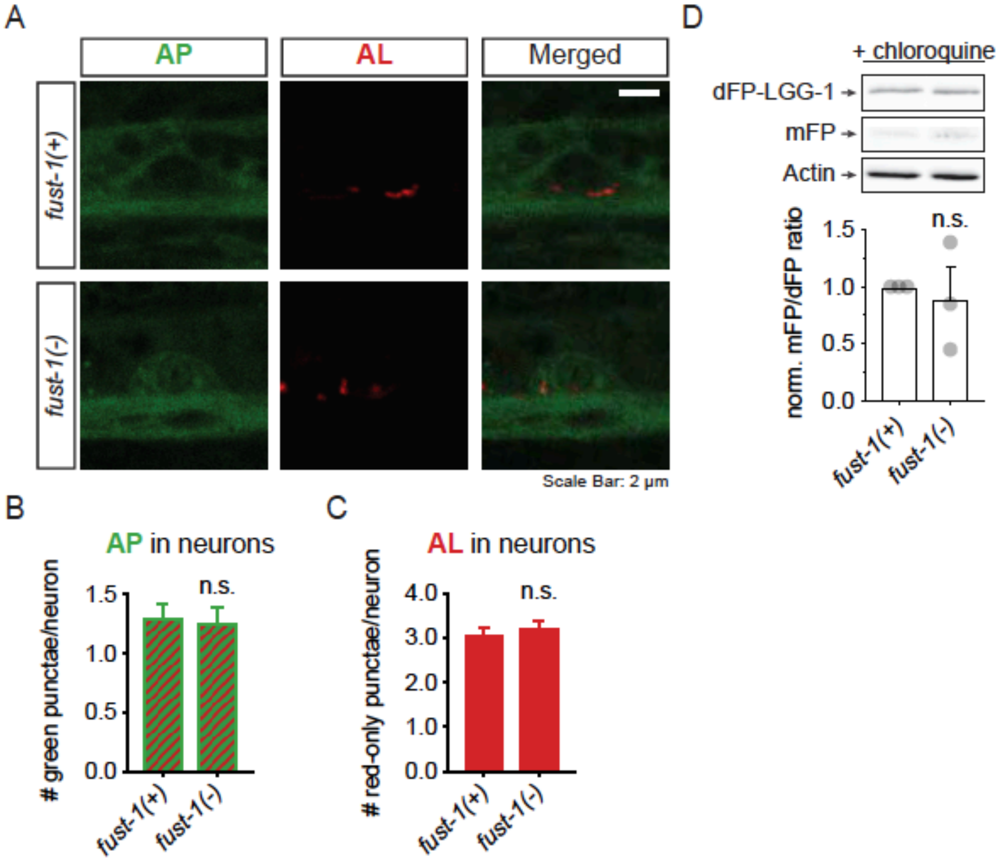


**Figure S5. Loss of *fust-1* does not disrupt autophagy in motor neurons (related to Figure 3).**

**(A)** Representative images of *fust-1* loss of function animals and controls on day 1 of adulthood expressing mCherry::GFP::LGG-1 reporter in motor neurons. Scale Bar = 2 um.

**(B)** Quantification of green+red APs in the motor neurons of *fust-1* loss of function animals and controls. Loss of *fust-1* had no impact on the number of APs per neuron compared to non-transgenic *fust-1(+)* N2 animals. N > 117 neurons per genotype. Data collected from three independent trials. Error bars indicate SEM. Pairwise t-test.

**(C)** Quantification of red-only ALs in the motor neurons of *fust-1* loss of function animals and controls. Loss of *fust-1* had no impact on the number of APs per neuron compared to non-transgenic *fust-1(+)* N2 animals. N > 117 neurons per genotype. Data collected from three independent trials. Error bars indicate SEM. Pairwise t-test.

**(D)** Neuronal autophagy was further examined by Western Blot using a different LGG-1 transgene double-tagged with two mono-fluorescent (mFP) proteins. Top: LGG-1 double fluorescent protein (dFP::LGG-1) used in this study. Lysosomal hydrolases cleave a flexible linker sequence between two mono-fluorescent proteins (mFP) fused to the LGG-1 reporter. Measurement of mFP to dFP::LGG-1 protein levels by Western Blot permits the quantitation of neuronal autophagy [36]. Middle: representative Western Blot images of dFP::LGG-1, mFP and actin protein bands from *fust-1* loss of function animals and *fust-1(+)* N2 controls, after overnight treatment with chloroquine. Chloroquine inhibits lysosomal activity [36], thus allowing for the visualization of neuronal mFP proteins in autolysosomes by Western Blot. Bottom: quantitation of neuronal mFP/dFP::LGG-1 protein ratio in *fust-1(-)* loss of function animals normalized against *fust-1(+)* controls. Loss of *fust-1* has no impact on mFP/dFP::LGG-1 protein ratio compared to *fust-1(+)* controls. Three independent trials. Error bars indicate -± SEM. Pairwise t-test. *fust-1(-): fust-1(tm4439)*.

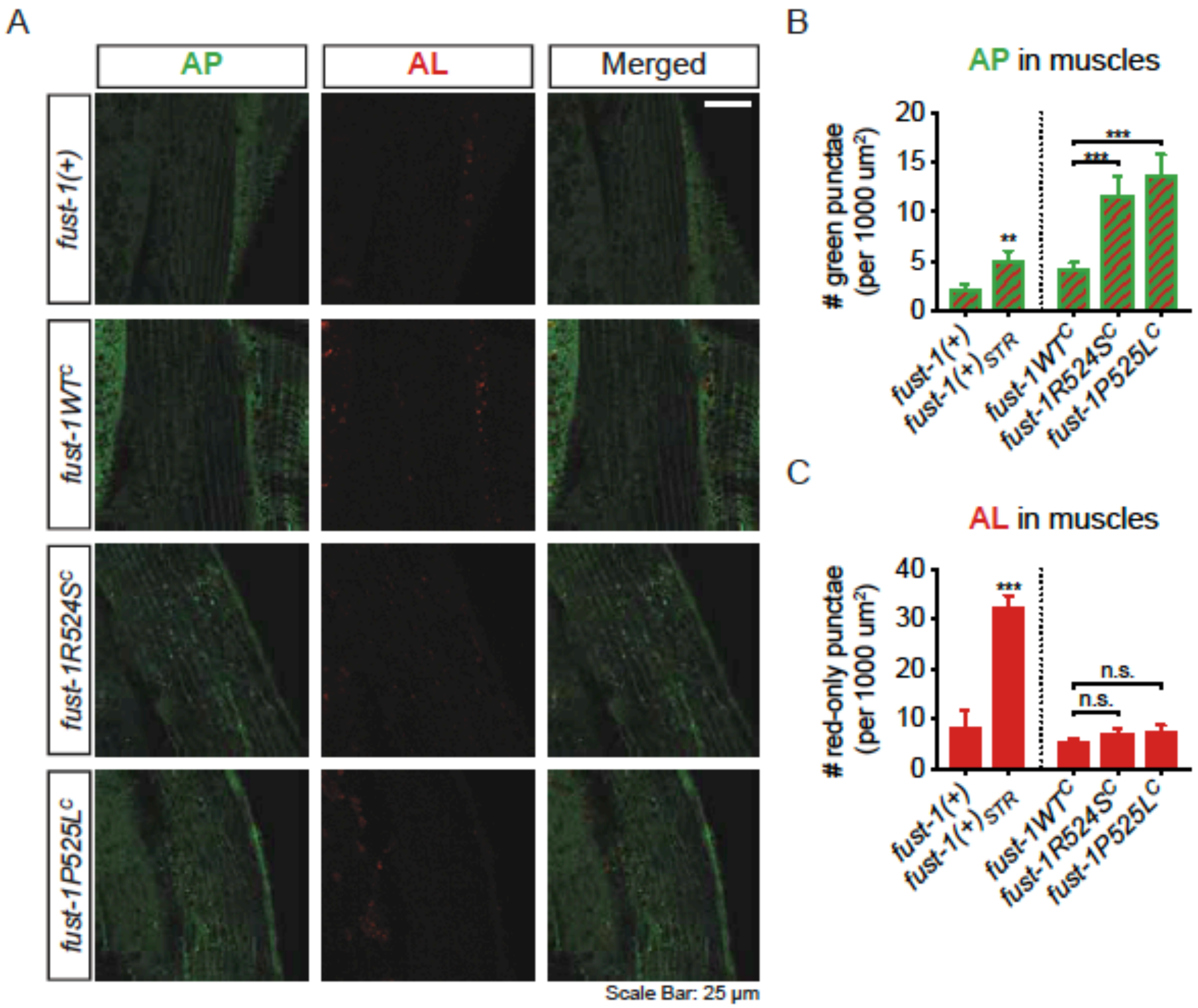


**Figure S6. ALS *fust-1* alleles disrupts autophagy in muscles (related to Figure 3).**

**(A)** Representative images of ALS *fust-1* animals and controls expressing the mCherry::GFP::LGG-1 reporter in muscles. Animals on day 1 of adulthood were mechanically paralyzed with microbeads to monitor autophagy in muscles. Scale bar = 25 um.

**(B)** Quantification of green+red APs in the muscles of ALS *fust-1* animals and controls. As a control, we examined autophagy in starved animals. Starvation increased the number of green+red AP punctae in motor neurons compared to satiated non-transgenic *fust-1(+)* animals. ALS *fust-1* alleles increased the number of APs per neuron compared to *fust-1WT*$^{C}$ controls. N > 31 animals per genotype. Data collected from three independent trials. Error bars indicate SEM. Pairwise t-test: * $p < 0.05$; ** $p < 0.01$; *** $p < 0.001$. *fust-1(+)*$_{STR}$: overnight starved animals.

**(C)** Quantification of red-only ALs in the muscles of ALS *fust-1* animals and controls. As a control, we examined autophagy in starved non-transgenic *fust-1(+)* animals. Starvation increased the number of red-only AL punctae in motor neurons compared to satiated non-transgenic *fust-1(+)* animals. By contrast, ALS *fust-1* alleles had no impact on the number ALs per neuron compared to *fust-1WT*$^{C}$ controls. N > 31 animals per genotype. Data collected from three independent trials. Error bars indicate SEM. Pairwise t-test: * $p < 0.05$; ** $p < 0.01$; *** $p < 0.001$. *fust-1(+)*$_{STR}$: overnight starved animals.

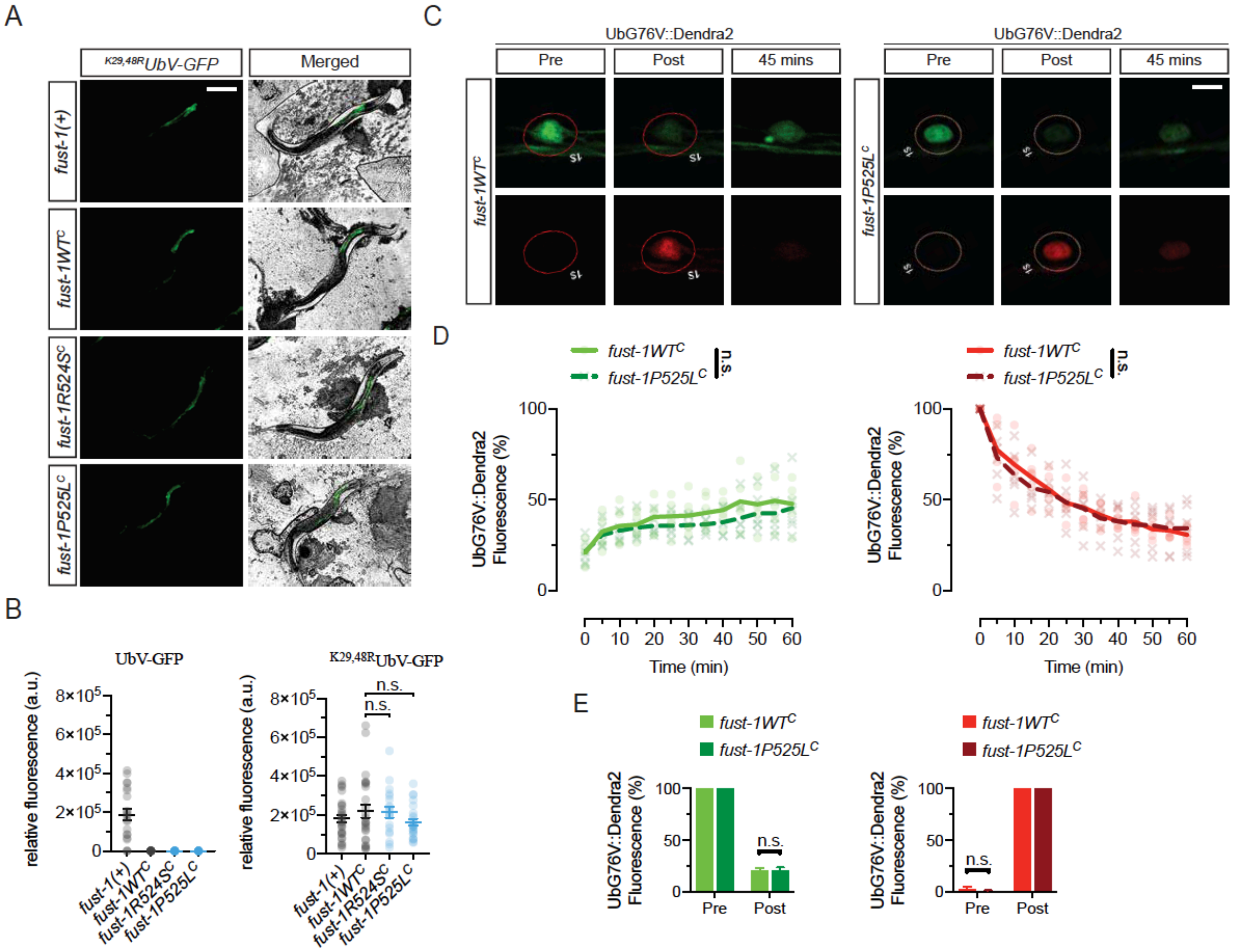


**Figure S7. ALS *fust-1* alleles have no impact on the ubiquitin proteasome system (UPS) (related to Figure 3).**

**(A)** Ubiquitin proteasome system (UPS) activity was examined in vivo by monitoring the levels of two different ubiquitin-GFP fusion substrates broadly expressed under the *sur-5* promoter: a non-cleavable ubiquitin reporter (UbV) fused with GFP (UbV-GFP), and a K29,48R mutant ubiquitin reporter fused with GFP ($^{K29,48R}$UbV-GFP) [54]. Mutation of ubiquitin lysine (K) residues 29 and 48 with arginine (K29,48R) stabilizes the protein, likely by preventing the polyubiquitination of the substrate at K29 and K48 [54]. Representative images show the $^{K29, 48R}$UbV-GFP substrate in ALS *fust-1* animals and controls. Scale Bar = 150 um.

**(B)** Quantification of the whole animal fluorescence in ALS *fust-1* animals and controls expressing the UbV-GFP or $^{K29,48R}$UbV-GFP substrates. No difference between ALS *fust-1* animals and *fust-1WT$^{C}$* controls was detected for either one of the ubiquitin substrates. N > 21 animals per genotype. Data collected from two independent trials. Error bars indicate SEM. Pairwise t-test.

**(C)** UPS activity was examined in transgenic animals expressing a non-cleavable ubiquitin moiety (UbG76V) fused with a photoconvertible fluorescent tag (Dendra2) under the *F25B3.3* neuronal promoter [55]. Motor neurons expressing the green fluorescent UbG76V::Dendra2 substrate were photo converted to a red fluorescent state by 405 nm light excitation. After photoconversion, quantification of the green and red fluorescence signals allows for the determination of protein synthesis and degradation, respectively. Representative images show motor neurons in *fust-1P525L$^{C}$* animals and *fust-1WT$^{C}$* controls before (pre) and immediately after photoconversion (post), as well as 45 minutes after photoconversion. Scale Bar = 2 um.

**(D)** Quantification of green (left) and red (right) UbG76V::Dendra2 fluorescence in motor neurons of *fust-1P525L$^{C}$* animals and *fust-1WT$^{C}$* controls after photoconversion. Relative green and red UbG76V::Dendra2 fluorescence intensities over time were normalized against the green and red UbG76V::Dendra2 fluorescence intensities observed immediately before and after photoconversion, respectively. N = 6 motor neurons per genotype. Two-way repeated measures ANOVA. No main effects of genotype were observed (green fluorescence: $F_{(1, 10)} = 0.9729$, $p = 0.3472$; red fluorescence: $F_{(1, 10)} = 0.06308$, $p = 0.8068$).

**(E)** Relative conversion rate of green UbG76V::Dendra2 fluorescence (left) and into red UbG76V::Dendra2 fluorescence (right) was similar in motor neurons of *fust-1WT*$^{C}$ and *fust-1P525L*$^{C}$ animals. Error bars indicate SEM. Pairwise t-test.

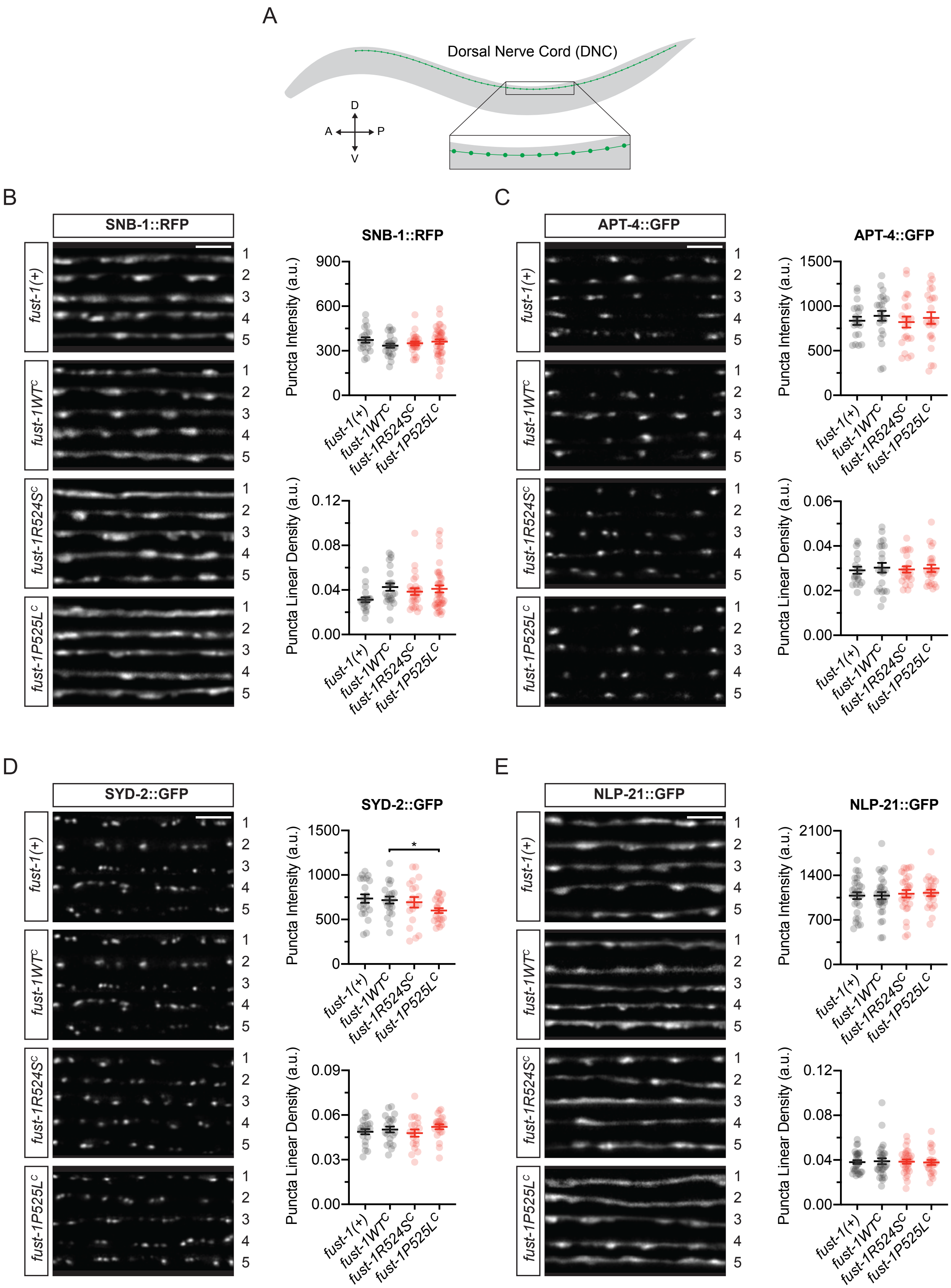


**Figure S8. ALS *fust-1* animals have normal levels and distribution of presynaptic proteins in motor neurons**

**(related to Figure 4).**

**(A)** Neuromuscular junction structures along the dorsal nerve cord were examined by measuring levels and distribution of fluorescently-labelled presynaptic proteins expressed in DA and DB cholinergic motor neurons.

**(B-E)** Representative images show fluorescently-labelled presynaptic proteins from 5 different animals for each genotype and marker. Graphs show quantification of levels (puncta intensity) and distribution (puncta linear density) of different fluorescently-labelled presynaptic proteins in ALS *fust-1* animals and controls. Levels and distribution of fluorescently-labelled presynaptic proteins SNB-1::RFP, APT-4::GFP and NLP-21::GFP, marking synaptic, endocytic and dense core vesicles, respectively, were normal in ALS *fust-1* animals. Levels, but not distribution, of SYD-2::GFP, an active zone marker, was slightly decreased in ALS *fust-1P525L*$^{C}$ animals compared to *fust-1WT*$^{C}$ controls. Both levels and distribution of SYD-2::GFP are normal in *fust-1R524S*$^{C}$ animals. N > 19 animals per genotype. Data were collected from at least three independent trials for each synaptic marker. Error bars indicate SEM. Pairwise t-test: * $p < 0.05$; ** $p < 0.01$; *** $p < 0.001$. Scale bar = 2 um.

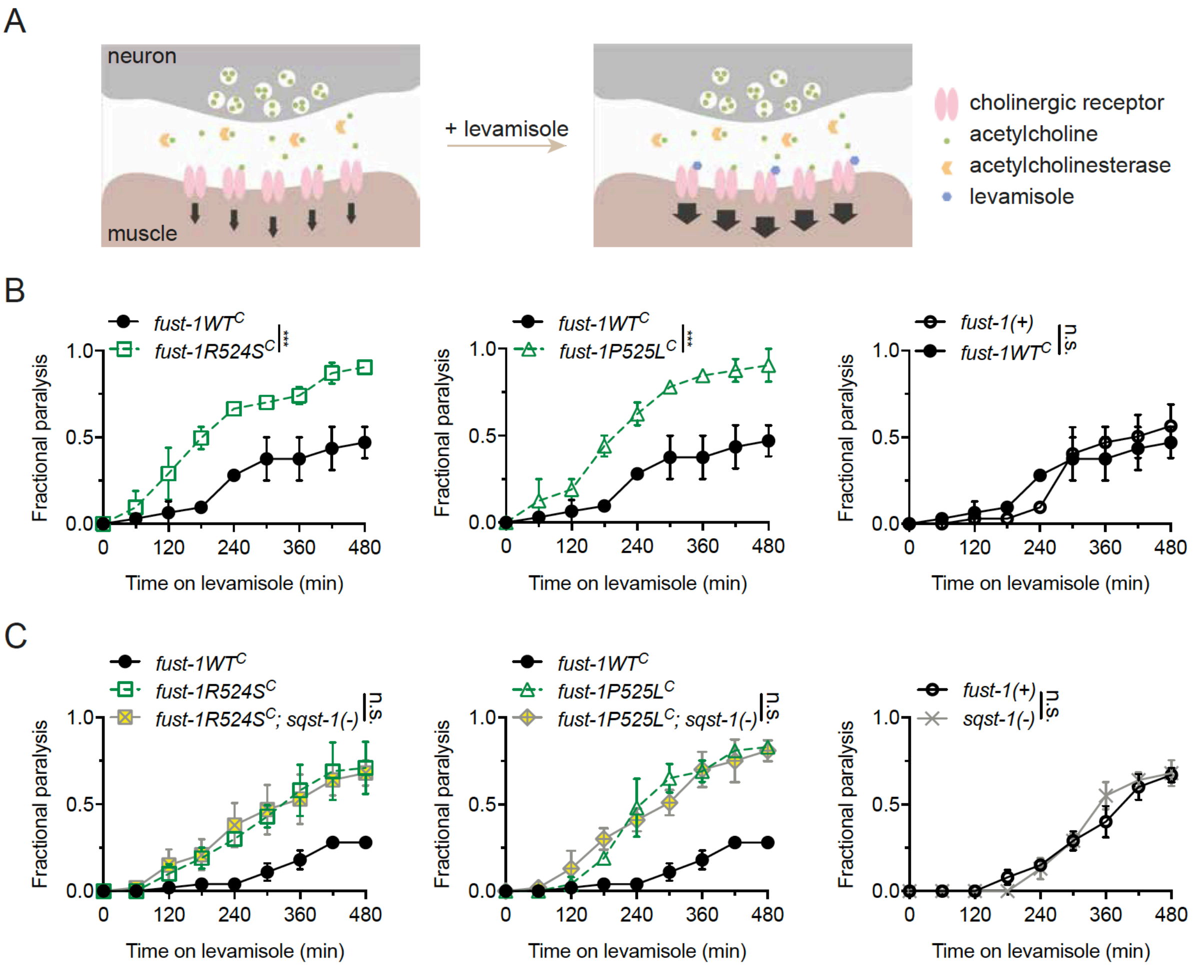


**Figure S9. ALS *fust-1* alleles disrupt muscular function. Loss of *sqst-1*, *C. elegans* ortholog for mammalian SQSTM1, does not suppress muscular function defects in ALS *fust-1* animals (related to Figure 4).**

**(A)** Muscular function was assessed with levamisole, a cholinergic receptor agonist. Activation of postsynaptic cholinergic receptors with levamisole leads to hyperexcitation of postsynaptic muscles, and paralysis. While wild type animals on levamisole paralyze over a characteristic time-course, hypersensitivity or resistance to levamisole suggests impaired muscular function.

**(B)** ALS *fust-1* animals are hypersensitive to levamisole compared to *fust-1WT$^C$* controls. N > 45 animals per genotype. Three independent trials. Error bars indicate SEM. Log-rank test: * $p < 0.05$; ** $p < 0.01$; *** $p < 0.001$.

**(C)** Loss of function allele *sqst-1(ok2892)* in ALS *fust-1* animals did not suppress levamisole sensitivity. N > 45 animals per genotype. Three independent trials. Error bars indicate SEM. Log-rank test: * $p < 0.05$; ** $p < 0.01$; *** $p < 0.001$.

A

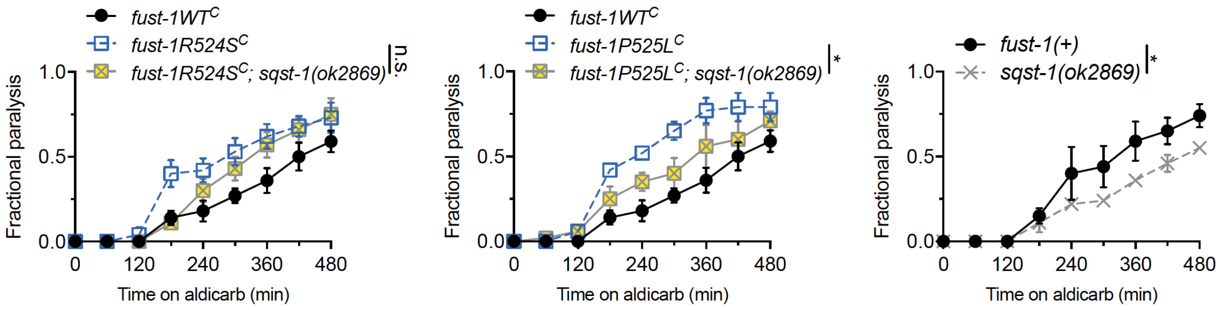


B

| | *fust-1R524S^C* | *lof; fust-1R524S^C* | *p*-value | *fust-1P525L^C* | *lof; fust-1P525L^C* | *p*-value |
|---|---|---|---|---|---|---|
| *daao-1* | 0.85 ± 0.02 | 0.73 ± 0.08 | 0.2193 | 0.81 ± 0.03 | 0.72 ± 0.10 | 0.4373 |
| *ubql-1* | 0.81 ± 0.07 | 0.77 ± 0.11 | 0.7743 | 0.71 ± 0.11 | 0.54 ± 0.03 | 0.2102 |
| *C34B7.2* | 0.78 ± 0.02 | 0.86 ± 0.01 | 0.0232 | 0.76 ± 0.06 | 0.79 ± 0.79 | 0.6779 |
| *djr-1.2* | 0.81 ± 0.00 | 0.75 ± 0.09 | 0.5415 | 0.71 ± 0.03 | 0.88 ± 0.07 | 0.0894 |

**Figure S10. Neuromuscular defects in double-mutant ALS *fust-1* animals (related to Figure 4).**

**(A)** A second *sqst-1(ok2869)* loss of function allele partially suppressed aldicarb hypersensitivity in *fust-1P525L*$^{C}$ animals. *sqst-1(ok2869)* allele causes a frameshift mutation in the second exon of *sqst-1*. N > 45 per genotype. Three independent trials. Log-rank test: * $p < 0.05$.

**(B)** Loss of function (lof) mutations in four other ALS genes in *C. elegans* do not suppress aldicarb hypersensitivity in ALS *fust-1* animals. Numbers indicate the average fraction of paralyzed animals after 8 hours of aldicarb treatment and SEM. N > 45 per genotype. Three independent trials. Pairwise t-test.

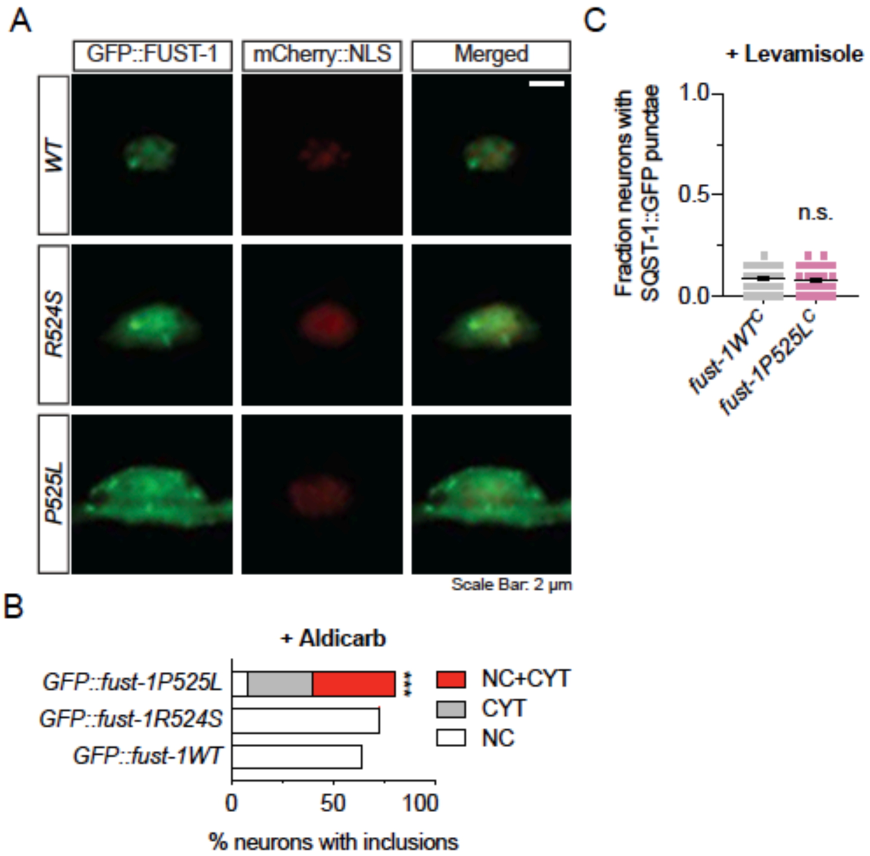


**Figure S11. Aldicarb induces stress in *C. elegans* (related to Figure 6).**

**(A)** Representative images of wild type and mutant GFP::FUST-1 in motor neurons after aldicarb treatment. Wild type GFP::FUST-1 formed nuclear inclusions in motor neurons after aldicarb treatment. GFP::FUST-1P525L formed both nuclear and cytoplasmic inclusions. GFP::FUST-1R524S formed mostly nuclear inclusions, with some cytoplasmic inclusions.

**(B)** Quantification of nuclear and cytoplasmic inclusions in the motor neurons of transgenic animals expressing wild type and mutant GFP::FUST-1. Data collected from three independent trials. N > 140 neurons per genotype. Chi-square test: * $p < 0.05$; ** $p < 0.01$; *** $p < 0.001$.

**(C)** 0.4 mM levamisole treatment for 4 hours does not induce SQST-1 accumulation in *C. elegans* motor neurons. N = 40 animals per genotype. Data collected from two independent trials. Pairwise t-test.

**Table S1. Primers**

| Primer Name | Sequence | Note |
|---|---|---|
| daao-1(tm3673)_geno_f | tttctgaagtcagctcaatg | |
| daao-1(tm3673)_geno_rint | taaattttcgttcgggtgg | |
| daao-1(tm3673)_geno_rext | tattccgaatctgaagggg | |
| sqst-1_geno_f | tgtcaacggaccaatctttg | |
| sqst-1_geno_r | cttcttgttcaagacgggct | |
| C34B7.2(tm5202)_geno_f | agcaacgatttcagatgg | |
| C34B7.2(tm5202)_geno_rint | caacaaatcgtacacatcc | |
| C34B7.2(tm5202)_geno_rext | aaattcagcaccaacgag | |
| ubql-1(tm1574)_geno_f | ttggaaggcaaattgattggag | Genotyping primers |
| ubql-1(tm1574)_geno_rint | cggagacggaaaagaacac | |
| ubql-1(tm1574)_geno_rext | cctgaagattcctaatagcttg | |
| djr-1.2(tm1496)_geno_f | ggtccctccagtgtttcgac | |
| djr-1.2(tm1496)_geno_r | tcaccgaataccgagcgtca | |
| fust-1(tm4439)_geno_f | agactacacgcgacgcacaac | |
| fust-1(tm4439)_geno_rint | ccggatccttggccgtatgg | |
| fust-1(tm4439)_geno_rext | tccgccctgtctgcaaaacatc | |
| fust-1_scr_f | gagtaggtggcggaaatg | |
| fust-1_scr_r | ggaagaatgagaggtggg | |
| unc-17p_f | taccactagttcccatccgcttcatccaaaagcggcgttc | |
| unc-17p_r | ttctcctttactcatctctctctctccccctggaatattttattttttc | |
| GFP_f | gggggagagagagagatgagtaaaggagaagaattgttcactggag | |
| GFP_r | cgatgctcctgaggctcccgatgctcccttgtagagctcgtccattccgtgggt<br>gattccggcggcgg | *Punc-17::GFP::fust-1::unc-54 3' UTR* plasmids cloning primers |
| fust-1_ex1_f | cgggagcctcaggagcatcgatgggttagtttctttttaatagtcgttttcttattg<br>g | |
| fust-1WT_ex8_r | tgaagagtaattggatcaatatggacggtaacgatcgccacccgg | |
| unc-54_3UTR_fust-1WT_f | taccgtccatattgatccaattactcttcaacatccctacatgctc | |
| unc-54_r | atgcggtacctaaaaattgcggtcataaactgaaacgtaac | |
| fust-1R524S_ex8_r | tgaagagtaattggatcaatatggagagtaacgatcgccacccgg | |
| unc-54_3UTR_fust-1R524S_f | tactctccatattgatccaattactcttcaacatccctacatgctc | |
| fust-1P525L_ex8_r | tgaagagtaattggatcaataaagacggtaacgatcgccacccgg | *fust-1* guide RNA plasmid cloning primers |
| unc-54_3UTR_fust-1P525L_f | taccgtctttattgatccaattactcttcaacatccctacatgctc | |
| fust-1_guide_f | cggtcacctccgttttagagctagaaatagc | |
| fust-1_guide_r | ataccgtccaaacatttagatttgcaattc | |
| fust-1WT_ssODN | gttaaaatttatggggagacgagttgatggctcaatatggacggtaacgatc<br>gccacccggtggtcctcctcgtcgctcgccaccaccacctccggatc | |
| fust-1R524S_ssODN | gttaaaatttatggggagacgagttgatggctcaatatggagagtaacgatc<br>gccacccggtggtcctcctcgtcgctcgccaccaccacctccggatc | Homologous recombination template primers |
| fust-1P525L_ssODN | gttaaaatttatggggagacgagttgatggctcaataaagacggtaacgatc<br>gccacccggtggtcctcctcgtcgctcgccaccaccacctccggatc | |

### Table S2. Strains

| Genotype |
| --- |
| fust-1(+) |
| rtEx960 [unc-17p::GFP::fust-1WT::unc-54 3'UTR]; otIs544 [cho-1(fosmid)::SL2::mCherry::H2B + pha-1(+)] |
| rtEx961 [unc-17p::GFP::fust-1R524S::unc-54 3'UTR]; otIs544 [cho-1(fosmid)::SL2::mCherry::H2B + pha-1(+)] |
| rtEx962 [unc-17p::GFP::fust-1P525L::unc-54 3'UTR]; otIs544 [cho-1(fosmid)::SL2::mCherry::H2B + pha-1(+)] |
| pha-1(+) III; fust-1(rt255 [fust-1WT$^{C}$ ]) II |
| pha-1(+) III; fust-1(rt256 [fust-1R524S$^{C}$ ]) II |
| pha-1(+) III; fust-1(rt255 [fust-1P525L$^{C}$ ]) II |
| fust-1(tm4439) II |
| sqst-1(ok2869) IV |
| sqst-1(ok2892) IV |
| fust-1(rt255 [fust-1WT$^{C}$ ]) II; nuIs175 [myo-2p::RFP, unc-129p::RFP::snb-1] X |
| fust-1(rt256 [fust-1R524S$^{C}$ ]) II; nuIs175 [myo-2p::RFP, unc-129p::RFP::snb-1] X |
| fust-1(rt255 [fust-1P525L$^{C}$ ]) II; nuIs175 [myo-2p::RFP, unc-129p::RFP::snb-1] X |
| nuIs175 [myo-2p::RFP, unc-129p::RFP::snb-1] X |
| fust-1(rt255 [fust-1WT$^{C}$ ]) II; nuIs160 [unc-129p::GFP::syd-2] III |
| fust-1(rt256 [fust-1R524S$^{C}$ ]) II; nuIs160 [unc-129p::GFP::syd-2] III |
| fust-1(rt255 [fust-1P525L$^{C}$ ]) II; nuIs160 [unc-129p::GFP::syd-2] III |
| nuIs160 [unc-129p::GFP::syd-2] III |
| fust-1(rt255 [fust-1WT$^{C}$ ]) II; nuIs184 [myo-2p::GFP, unc-129p::apt-4::GFP] X |
| fust-1(rt256 [fust-1R524S$^{C}$ ]) II; nuIs184 [myo-2p::GFP, unc-129p::apt-4::GFP] X |
| fust-1(rt255 [fust-1P525L$^{C}$ ]) II; nuIs184 [myo-2p::GFP, unc-129p::apt-4::GFP] X |
| nuIs184 [myo-2p::GFP, unc-129p::apt-4::GFP] X |
| fust-1(rt255 [fust-1WT$^{C}$ ]) II; nuIs183[unc-129p::venus::nlp-21] III |
| fust-1(rt256 [fust-1R524S$^{C}$ ]) II; nuIs183[unc-129p::venus::nlp-21] III |
| fust-1(rt255 [fust-1P525L$^{C}$ ]) II; nuIs183[unc-129p::venus::nlp-21] III |
| nuIs183 [unc-129p::venus::nlp-21] III |
| rtIs97 [rab-3p::Cerulean::Venus::lgg-1] |
| fust-1(tm4439) II; rtIs97 [rab-3p::Cerulean::Venus::lgg-1] |
| fust-1(rt255 [fust-1WT$^{C}$ ]) II; rtIs97 [rab-3p::Cerulean::Venus::lgg-1] |
| fust-1(rt256 [fust-1R524S$^{C}$ ]) II; rtIs97 [rab-3p::Cerulean::Venus::lgg-1] |
| fust-1(rt255 [fust-1P525L$^{C}$ ]) II; rtIs97 [rab-3p::Cerulean::Venus::lgg-1] |
| sqIs11[ rol-6, lgg-1p::mCherry::GFP::lgg-1] |
| atg-7(bp411); sqIs11 [ rol-6, lgg-1p::mCherry::GFP::lgg-1] |
| fust-1(tm4439); sqIs11 [ rol-6, lgg-1p::mCherry::GFP::lgg-1] |
| fust-1(rt255 [fust-1WT$^{C}$ ]) II; sqIs11 [ rol-6, lgg-1p::mCherry::GFP::lgg-1] |
| fust-1(rt256 [fust-1R524S$^{C}$ ]) II; sqIs11 [ rol-6, lgg-1p::mCherry::GFP::lgg-1] |
| fust-1(rt255 [fust-1P525L$^{C}$ ]) II; sqIs11 [ rol-6, lgg-1p::mCherry::GFP::lgg-1] |
| [sur-5p::UbV::GFP] |
| fust-1(rt255 [fust-1WT$^{C}$ ]) II; [sur-5p::UbV::GFP] |
| fust-1(rt256 [fust-1R524S$^{C}$ ]) II; [sur-5p::UbV::GFP] |
| fust-1(rt255 [fust-1P525L$^{C}$ ]) II; [sur-5p::UbV::GFP] |
| [sur-5p::K29,48RUbV::GFP] |
| fust-1(rt255 [fust-1WT$^{C}$ ]) II; [sur-5p::K29,48RUbV::GFP] |
| fust-1(rt256 [fust-1R524S$^{C}$ ]) II; [sur-5p::K29,48RUbV::GFP] |
| fust-1(rt255 [fust-1P525L$^{C}$ ]) II; [sur-5p::K29,48RUbV::GFP] |
| fust-1(rt255 [fust-1WT$^{C}$ ]) II; xzEx12 [PF25B3.3::UbG76V::Dendra2] |
| fust-1(rt255 [fust-1P525L$^{C}$ ]) II; xzEx12 [PF25B3.3::UbG76V::Dendra2] |
| fust-1(rt255 [fust-1WT$^{C}$ ]) II; otIs544 [cho-1(fosmid)::SL2::mCherry::H2B + pha-1(+)] |
| fust-1(rt256 [fust-1R524S$^{C}$ ]) II; otIs544 [cho-1(fosmid)::SL2::mCherry::H2B + pha-1(+)] |
| fust-1(rt255 [fust-1P525L$^{C}$ ]) II; otIs544 [cho-1(fosmid)::SL2::mCherry::H2B + pha-1(+)] |
| rtEx960 [unc-17p::GFP::fust-1WT::unc-54 3'UTR]; sqst-1(ok2892) IV; otIs544 [cho-1(fosmid)::SL2::mCherry::H2B + pha-1(+)] |
| rtEx961 [unc-17p::GFP::fust-1R524S::unc-54 3'UTR]; sqst-1(ok2892) IV; otIs544 [cho-1(fosmid)::SL2::mCherry::H2B + pha-1(+)] |
| rtEx962 [unc-17p::GFP::fust-1P525L::unc-54 3'UTR]; sqst-1(ok2892) IV; otIs544 [cho-1(fosmid)::SL2::mCherry::H2B + pha-1(+)] |
| sqst-1(ok2892) IV; sqIs11[ rol-6, lgg-1p::mCherry::GFP::lgg-1] |
| fust-1(rt255 [fust-1WT$^{C}$ ]) II; sqst-1(ok2892) IV; sqIs11 [ rol-6, lgg-1p::mCherry::GFP::lgg-1] |
| fust-1(rt256 [fust-1R524S$^{C}$ ]) II; sqst-1(ok2892) IV; sqIs11 [ rol-6, lgg-1p::mCherry::GFP::lgg-1] |
| fust-1(rt255 [fust-1P525L$^{C}$ ]) II; sqst-1(ok2892) IV; sqIs11 [ rol-6, lgg-1p::mCherry::GFP::lgg-1] |
| bpIs151 [unc-76(+), sqst-1p::sqst-1::GFP] |
| fust-1(rt255 [fust-1WT$^{C}$ ]) II; bpIs151 [unc-76(+),sqst-1p::sqst-1::GFP] |
| fust-1(rt256 [fust-1P525L$^{C}$ ]) II; bpIs151 [unc-76(+),sqst-1p::sqst-1::GFP] |
| fust-1(rt255 [fust-1WT$^{C}$ ]) II; sqst-1(ok2892) IV |
| fust-1(rt256 [fust-1R524S$^{C}$ ]) II; sqst-1(ok2892) IV |
| fust-1(rt255 [fust-1P525L$^{C}$ ]) II; sqst-1(ok2892) IV |
| fust-1(rt255 [fust-1WT$^{C}$ ]) II; sqst-1(ok2869) IV |
| fust-1(rt256 [fust-1R524S$^{C}$ ]) II; sqst-1(ok2869) IV |
| fust-1(rt255 [fust-1P525L$^{C}$ ]) II; sqst-1(ok2869) IV |
| daao-1(tm3673) IV |
| fust-1(rt256 [fust-1R524S$^{C}$ ]) II; daao-1(tm3673) IV |
| fust-1(rt255 [fust-1P525L$^{C}$ ]) II; daao-1(tm3673) IV |
| ubql-1(tm1574) I |
| fust-1(rt256 [fust-1R524S$^{C}$ ]) II; ubql-1(tm1574) I |
| fust-1(rt255 [fust-1P525L$^{C}$ ]) II; ubql-1(tm1574) I |
| djr-1.2(tm1496) V |
| fust-1(rt256 [fust-1R524S$^{C}$ ]) II; djr-1.2(tm1496) V |
| fust-1(rt255 [fust-1P525L$^{C}$ ]) II; djr-1.2(tm1496) V |
| C34B7.2(tm5202) I |
| fust-1(rt256 [fust-1R524S$^{C}$ ]) II; C34B7.2(tm5202) I |
| fust-1(rt255 [fust-1P525L$^{C}$ ]) II; C34B7.2(tm5202) I |
| daf-16(mu86); muIs109 [Pdaf-16::gfp::daf-16cDNA + Podr-1::rfp] |
| daf-16(mu86); muIs71 [daf-16ap::GFP::daf-16a(bKO)) + rol-6(su1006)] |
